# The Dense-Core Vesicle Maturation Protein CCCP-1 Binds RAB-2 and Membranes through its C-terminal Domain

**DOI:** 10.1101/105668

**Authors:** Jérôme Cattin-Ortolá, Irini Topalidou, Annie Dosey, Alexey J. Merz, Michael Ailion

**Affiliations:** Departments of Biochemistry, University of Washington, Seattle, WA, 98195 USA; Physiology and Biophysics, University of Washington, Seattle, WA, 98195 USA

**Author notes:** To whom correspondence should be addressed: Michael Ailion, Department of Biochemistry, University of Washington, Box 357350, 1705 NE Pacific St, Seattle, WA 98195.

**Keywords:** Membrane trafficking, dense-core vesicle, Rab, GTPase, *Caenorhabditis elegans*, insulinoma 832/13 cells, coiled-coil domain, golgin, lipid binding protein

## Abstract

Dense-core vesicles (DCVs) are secretory organelles that store and release modulatory neurotransmitters from neurons and endocrine cells. Recently, the conserved coiled-coil protein CCCP-1 was identified as a component of the DCV biogenesis pathway in the nematode *C. elegans*. CCCP-1 binds the small GTPase RAB-2 and colocalizes with it at the trans-Golgi. Here we report a structure-function analysis of CCCP-1 to identify domains of the protein important for its localization, binding to RAB-2, and function in DCV biogenesis. We find that the CCCP-1 C-terminal domain (CC3) has multiple activities. CC3 is necessary and sufficient for CCCP-1 localization and for binding to RAB-2, and is required for the function of CCCP-1 in DCV biogenesis. Additionally, CCCP-1 binds membranes directly through its CC3 domain, indicating that CC3 may comprise a previously uncharacterized lipid-binding motif. We conclude that CCCP-1 is a coiled-coil protein that binds an activated Rab and localizes to the Golgi via its C-terminus, properties similar to members of the golgin family of proteins. CCCP-1 also shares biophysical features with golgins; it has an elongated shape and forms oligomers.

**Synopsis statement:** CCCP-1 is a coiled-coil protein important for dense-core vesicle (DCV) biogenesis. A structure-function analysis of CCCP-1 shows that its C-terminal domain is required for (1) localization to membrane compartments near the trans-Golgi, (2) binding to activated RAB-2, (3) function in DCV biogenesis, and (4) direct binding to membranes. CCCP-1 has an elongated shape and forms oligomers. These findings suggest that CCCP-1 resembles members of the golgin family of proteins that act as membrane tethers.

## 1. INTRODUCTION

Dense-core vesicles (DCVs) are specialized organelles of neurons and endocrine cells. DCVs store and release modulatory peptides and biogenic amines, signaling molecules that control a multitude of processes including synaptic plasticity, feeding behavior, and glucose homeostasis^1–4^. However, DCV biogenesis is not well understood, especially at the molecular level. Immature DCVs bud off from the trans-Golgi and undergo several maturation steps, including homotypic fusion with other immature DCVs, vesicle acidification, peptide processing, and sorting of cargos^1–4^. In recent years, a small but growing number of proteins have been identified as playing roles in DCV biogenesis, beginning to shed some light on the molecular mechanisms underlying the steps in DCV maturation^5–18^. In one approach to identifying proteins important for DCV biogenesis, genetic screens were performed in *Caenorhabditis elegans*^19–25^. These screens identified several molecules important for early stages of DCV biogenesis and maturation, including the small G protein RAB-2 and several RAB-2 effectors, including the conserved coiled-coil protein CCCP-1^24^. Here we perform a structure-function analysis of CCCP-1 to better understand its biochemical activities and role in DCV biogenesis.

RAB-2 is a highly-conserved member of the Rab family of small G proteins. Rabs are membrane-associated proteins that function as molecular switches that alternate between a GTP-bound active state and a GDP-bound inactive state^26^. Activated Rabs recruit effectors that execute many aspects of membrane trafficking including vesicle transport and vesicle tethering^26^. CCCP-1 was recently identified as a potential RAB-2 effector^24,27^. In *C. elegans*, RAB-2 and CCCP-1 colocalize near the trans-Golgi and function in the same genetic pathway to regulate locomotion and DCV cargo sorting in neurons^24^. Though CCCP-1 is critical for proper DCV biogenesis, its specific molecular functions remain unclear. Here we conduct a structure-function analysis to determine which domains of CCCP-1 are important for its localization, binding to RAB-2, and function in DCV biogenesis. Our analysis shows that CCCP-1 has structural and functional features in common with the golgins, a family of tethers that also bind activated Rabs and regulate vesicular trafficking at the Golgi^28–32^. These findings suggest that CCCP-1 may act as a membrane tether.

## 2. RESULTS

### 2.1 CCCP-1 is localized near immature DCVs and the trans-Golgi via its C-terminal domain CC3

To assess the localization of full length CCCP-1 in *C. elegans* neurons, we generated transgenic worms expressing GFP-tagged CCCP-1. When expressed under its own promoter, either as a single-copy integration or as multiple copies in an extrachromosomal array, CCCP-1::GFP levels were too low to be detected in most transgenic lines. However, these constructs are functional since they rescued the locomotion defects of a *cccp-1* mutant (see below). Because CCCP-1 is expressed predominantly in *C. elegans* neurons^24^, we expressed CCCP-1::GFP under the stronger pan-neuronal *rab-3* promoter. CCCP-1 localized mainly to perinuclear puncta in the soma of ventral cord motor neurons (Figure 1A, top) but also localized to puncta in axons in the dorsal nerve cord (Figure 1A, bottom), suggesting that some CCCP-1 is transported out of the cell body.

**FIGURE 1.**
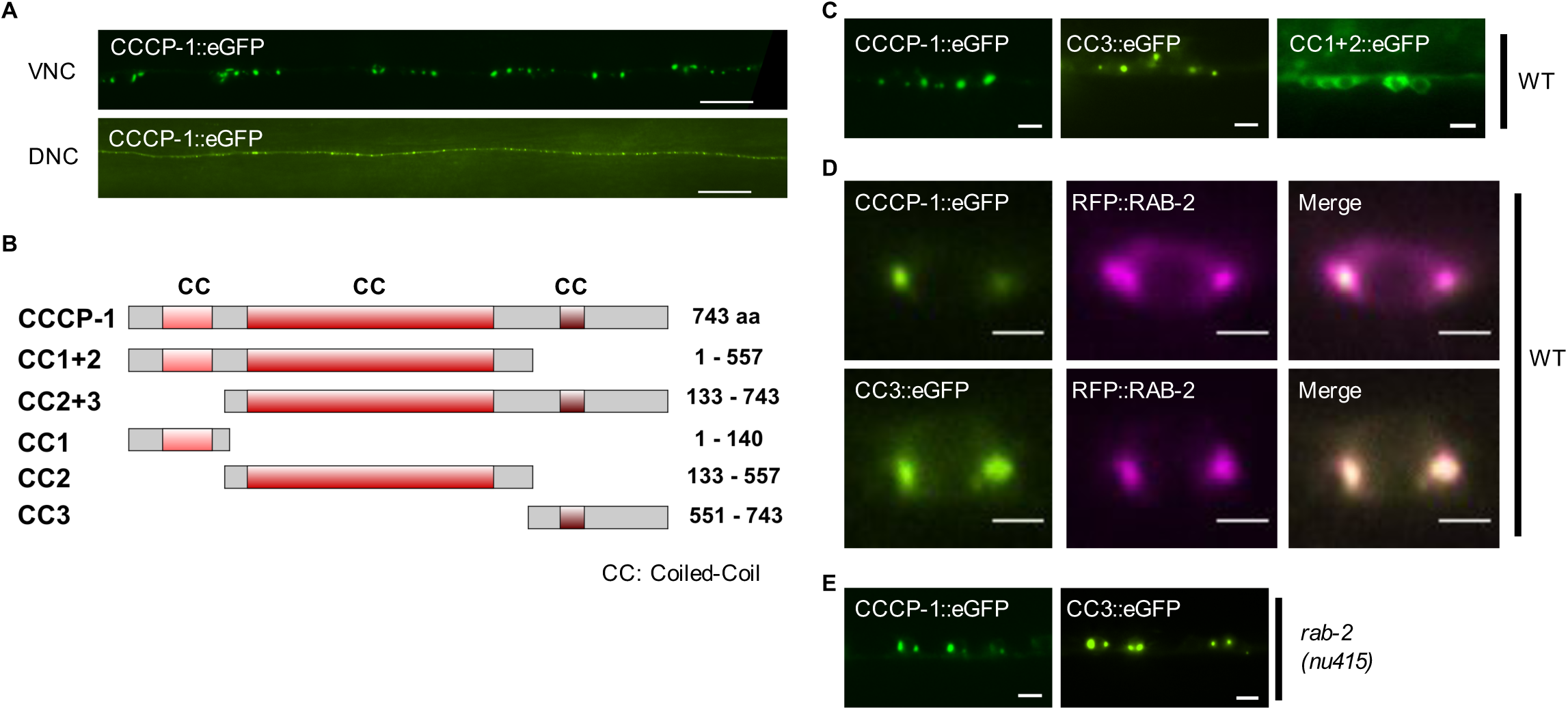
The CC3 C-terminal domain of CCCP-1 localizes the protein. A, CCCP-1 localizes to perinuclear and axonal puncta. Representative images of CCCP-1::eGFP expressed under the pan-neuronal *rab-3* promoter, showing cell bodies of *C. elegans* ventral cord motor neurons (VNC) and axons of the dorsal nerve cord (DNC). Scale bar: 20 µm. B, Domain structure of worm CCCP-1 and its different fragments. The CC1, CC2 and CC3 fragments carry the predicted coiled-coil domains. C, CC3 is necessary and sufficient for CCCP-1 localization. Representative images of the cell bodies of multiple adjacent ventral cord motor neurons expressing CCCP-1::eGFP, CC3::eGFP or CC1 + 2::eGFP, in wild-type animals. Scale bar: 5 µm. D, CC3 is necessary and sufficient for localization of the protein to RAB-2 positive membrane structures. Representative images of a single ventral cord motor neuron cell body coexpressing CCCP-1::eGFP or CC3::eGFP and tagRFP::RAB-2 in neurons of wild-type animals. Scale bar: 2 µm. E, CCCP-1 and CC3 localization does not require RAB-2. Representative images of ventral cord motor neuron cell bodies expressing CCCP-1::eGFP or CC3::eGFP in a *rab-2(nu415)* mutant background. Scale bar: 5 µm.

To determine which domain of CCCP-1 mediates its localization, we generated transgenic worm lines carrying different GFP-tagged CCCP-1 fragments (Figure 1B) expressed under the pan-neuronal *rab-3* promoter. Like full-length CCCP-1::GFP, a fragment containing the 193 C-terminal amino acids (CC3::GFP) localized to large perinuclear puncta (Figure 1C). Like CCCP-1::GFP, CC3::GFP had puncta in the dorsal nerve cord as well (data not shown). In contrast, CC1 + 2::GFP was localized diffusely in the cytoplasm (Figure 1C) as well as in the dorsal nerve cord (data not shown). These results are consistent with data showing that the N-terminal half of human CCCP1/CCDC186 localized to the cytosol and a C-terminal fragment larger than CC3 localized to perinuclear structures in COS cells^27^.

To test whether CC3 is sufficient for proper localization of CCCP-1, we performed colocalization experiments. We previously reported that CCCP-1 colocalized with the small GTPase RAB-2 near the trans-Golgi in *C. elegans* neurons^24^. Similar to full length CCCP-1::GFP, CC3::GFP colocalized with RFP::RAB-2 in *C. elegans* neurons (Figure 1D). Thus, the C-terminal domain CC3 is both necessary and sufficient for CCCP-1 localization.

To test whether CC3 is also necessary and sufficient for localization of mammalian CCCP1/CCDC186, we imaged GFP-tagged rat CCCP1/CCDC186 and CCCP1 fragments in rat insulinoma 832/13 cells^33^. These cells have secretory granules similar to neuronal DCVs and express the mammalian ortholog of CCCP-1, CCCP1/CCDC186^25^. Moreover, they are larger cells than *C. elegans* neurons and have a more easily visualized separation of different organelles.

In 832/13 cells, we found that endogenous CCCP1 localizes to perinuclear structures that overlapped with chromogranin A (CgA), a DCV cargo expected to localize to the trans-Golgi, immature DCVs, and mature DCVs (Figure 2A and 2B). CCCP1 localization was restricted to a perinuclear area adjacent to the cis-Golgi marker GM130 (Figure 2A). However, the CCCP1 signal overlaps more with CgA than with GM130 (Figure 2B). For technical reasons, we could not test the localization of endogenous CCCP1 with the trans-Golgi marker TGN38. CCCP1::GFP had a similar localization pattern to endogenous CCCP1. Like endogenous CCCP1, CCCP1::GFP mostly co-localized with CgA and was adjacent to GM130 (Figure 2C and 2D). CCCP1::GFP also partially colocalized with the trans-Golgi marker TGN38. Thus, as in *C. elegans* neurons and COS cells^24,27^, rat CCCP1 localizes near the trans-Golgi in 832/13 cells (Figure 2A-2D). Because no CCCP1 signal could be observed near the cell periphery and because CCCP1 localizes more closely with the DCV marker CgA than with TGN38 (Figure 2D), our results suggest that CCCP1 localizes near immature DCVs.

**FIGURE 2.**
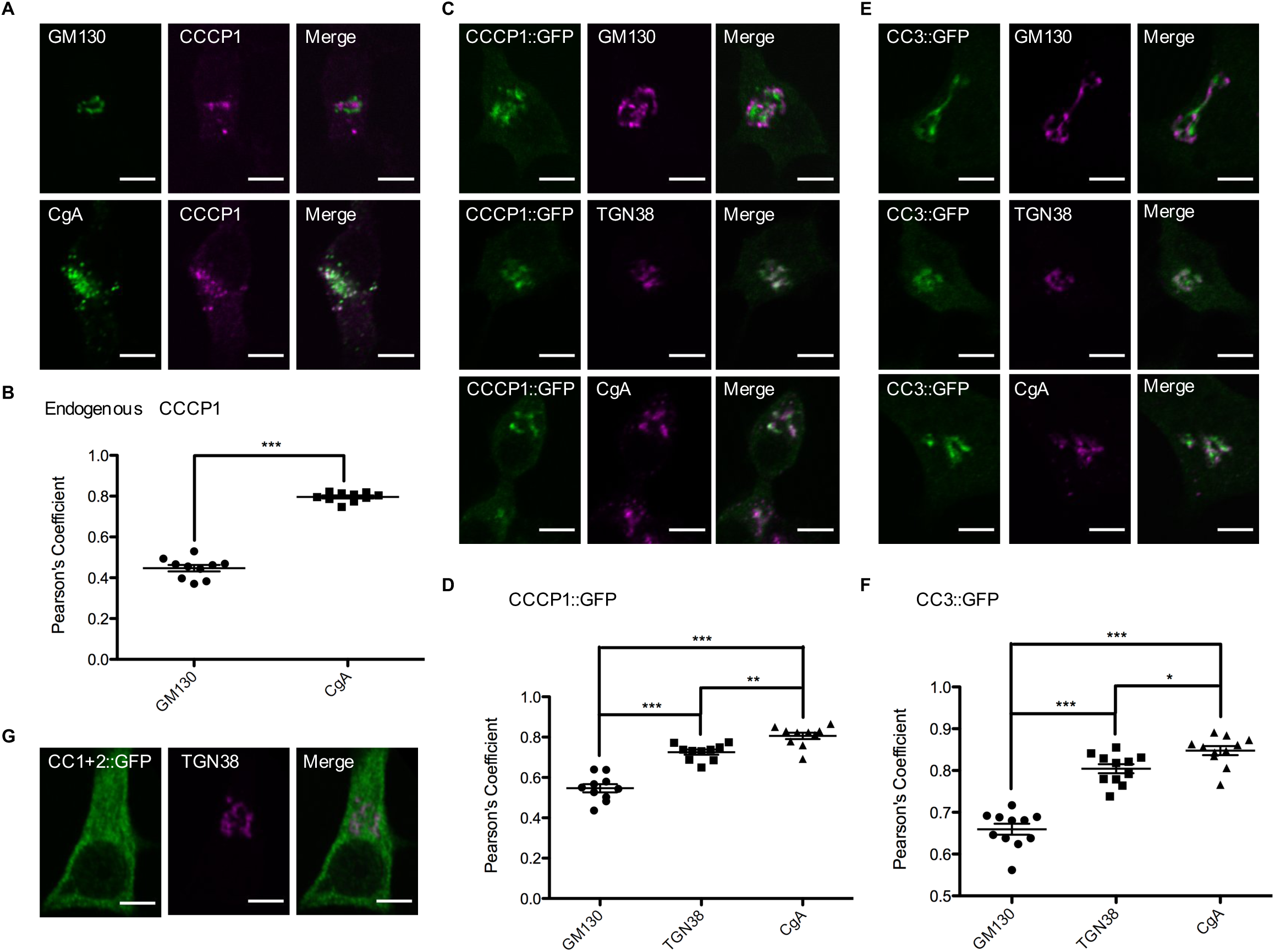
CC3 is necessary and sufficient for localization of rat CCCP1/CCDC186 near the trans-Golgi. A, Endogenous rat CCCP1 localizes to a perinuclear structure near a marker for DCV cargo, Chromogranin A (CgA). Representative images of 832/13 cells co-stained for endogenous CCCP1 and for the cis-Golgi marker GM130 (upper panel) and for the DCV marker CgA (lower panel). Scale bar: 5 µm. B, Quantification of endogenous CCCP1 colocalization with GM130 and CgA. ***: p < 0.001, n=10 each. C, CCCP1::GFP localizes to a perinuclear structure near the trans-Golgi marker TGN38 and the DCV marker CgA. Representative images of 832/13 cells co-stained for CCCP1::GFP and for the cis-Golgi marker GM130 (upper panel), the trans-Golgi marker TGN38 (middle panel) and for the DCV marker CgA (lower panel). Scale bar: 5 µm D, Quantification of CCCP1::GFP co-localization with GM130, TGN38 and CgA. ***: p < 0.001, **: p < 0.01, n=10 each. E, CC3::GFP localizes to a perinuclear structure near the trans-Golgi marker TGN38 and the DCV marker CgA. Representative images of 832/13 cells co-stained for CCCP1::GFP and for the cis-Golgi marker GM130 (upper panel), the trans-Golgi marker TGN38 (middle panel) and for the DCV marker ChgA (lower panel). Scale bar: 5 µm F, Quantification of CCCP1::GFP co-localization with GM130, TGN38 and ChgA. ***: p < 0.001, *: p < 0.05, n=11 each. G, CC1 + 2::GFP localizes to the cytoplasm. Representative images of 832/13 cells co-stained for CC1 + 2::GFP and for the trans-Golgi marker TGN38. Scale bar: 5 µm.

We tested the localization of CC3::GFP and found it similar to endogenous CCCP1 and CCCP1::GFP (Figure 2E and 2F). Finally, we found that, CC1 + 2::GFP localizes to the cytoplasm, as it does in *C. elegans* neurons (Figure 2G). Thus, CC3 is both necessary and sufficient for CCCP1/CCDC186 localization near immature DCVs and the trans-Golgi.

### 2.2 CCCP-1 binds via its C-terminal CC3 domain to RAB-2

Multiple lines of evidence suggest that CCCP-1 is an effector of the small GTPase RAB-2. CCCP-1 and RAB-2 act in the same genetic pathway in *C. elegans* and they colocalize in worm neurons and in COS cells^24,27^. In addition, yeast two-hybrid experiments suggested that CCCP-1 and RAB-2, from either worm or human, interact in a GTP-dependent manner^24,27^. The interaction of human CCCP1/CCDC186 with RAB2A was mapped to a C-terminal fragment of CCCP1/CCDC186^27^. To assess the RAB-2 interaction with CCCP-1 using purified *C. elegans* proteins, we performed GST pull-downs and found that CCCP-1 binds to RAB-2 loaded with the non-hydrolysable GTP analog guanosine 5’-O-[gamma-thio]-triphosphate (GTPγS, Figure 3, lane 1), but not to RAB-2 loaded with GDP (Figure 3, lane 2). The interaction of CCCP-1 with RAB-2 could be detected only by using an antibody to the His_6_ epitope tag, suggesting that the interaction is likely of moderate affinity.

**FIGURE 3.**
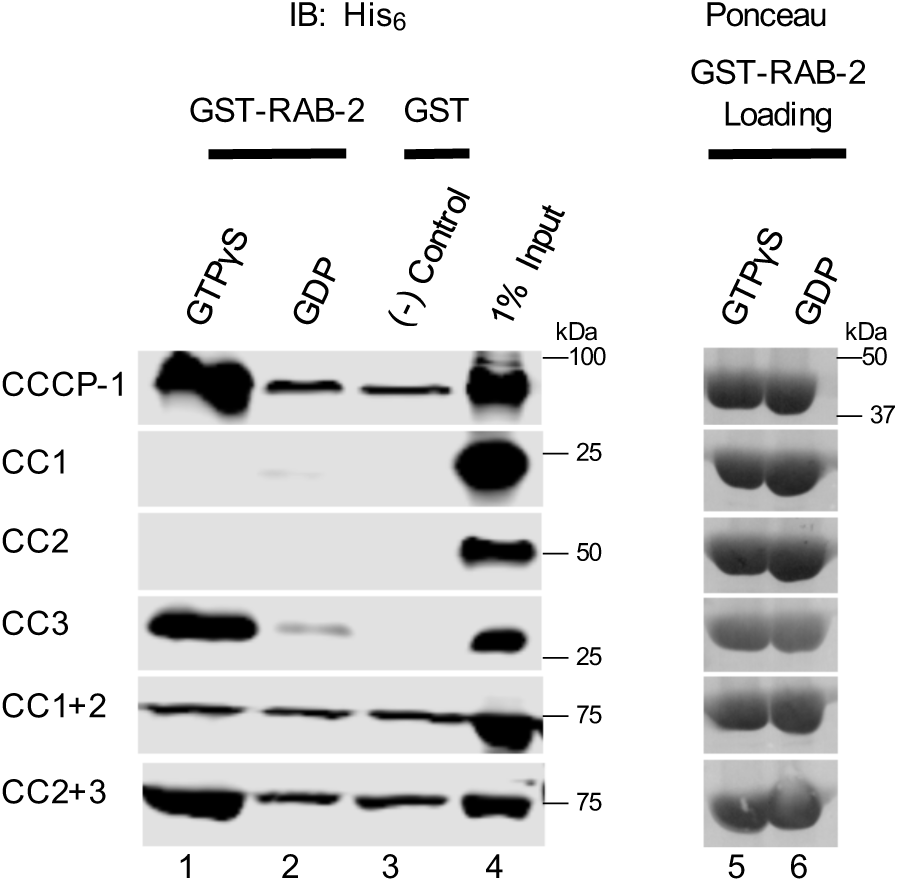
The CC3 domain of CCCP-1 binds directly to RAB-2. (Left) GST-pulldowns were performed using purified recombinant His_6_-CCCP-1 or its fragments and GST-RAB-2 loaded with either GTPγS (lane 1) or GDP (lane 2). GST on its own was used as a negative control (lane 3). 1% of the protein input is shown in lane 4. Samples were analyzed by Western blot against the His_6_ tag. (Right) GST-RAB-2 loading controls visualized by Ponceau staining show that approximately equal amounts of GST-RAB-2 were used (lanes 5 and 6). IB: immunoblot.

We also performed GST pull-down experiments using the CCCP-1 fragments. CC2 + 3 and CC3 bound to RAB-2 in a GTP-dependent manner but CC1 + 2 did not bind (Figure 3), demonstrating that CC3 is necessary and sufficient for RAB-2 binding.

In *C. elegans*, CCCP-1 localizes to perinuclear puncta in the absence of RAB-2^24^. This indicates that RAB-2 is not the sole determinant of CCCP-1 localization. We found that the localization of CC3 is also unaltered in the absence of RAB-2 (Figure 1E). Thus, the CC3 domain has at least two independent activities: RAB-2 binding and subcellular targeting.

### 2.3 CC3 is necessary for CCCP-1 function

To test whether CC3 is sufficient for the in vivo neuronal function of CCCP-1, we assayed CCCP-1 function in *C. elegans*. *cccp-1* mutant worms have a slow locomotion phenotype^24^. Because overexpression of CCCP-1 also causes slow locomotion, we instead made single-copy integrants of transgenes expressing the different CCCP-1 fragments under the endogenous *cccp-1* promoter and tested for rescue of the *cccp-1* mutant slow locomotion phenotype. Expression of full-length CCCP-1 rescued the locomotion defect of *cccp-1* mutants (Figure 4A), but neither CC1 + 2 nor CC3 rescued (Figure 4B). Thus, CC3 is necessary but not sufficient for CCCP-1 function.

**FIGURE 4.**
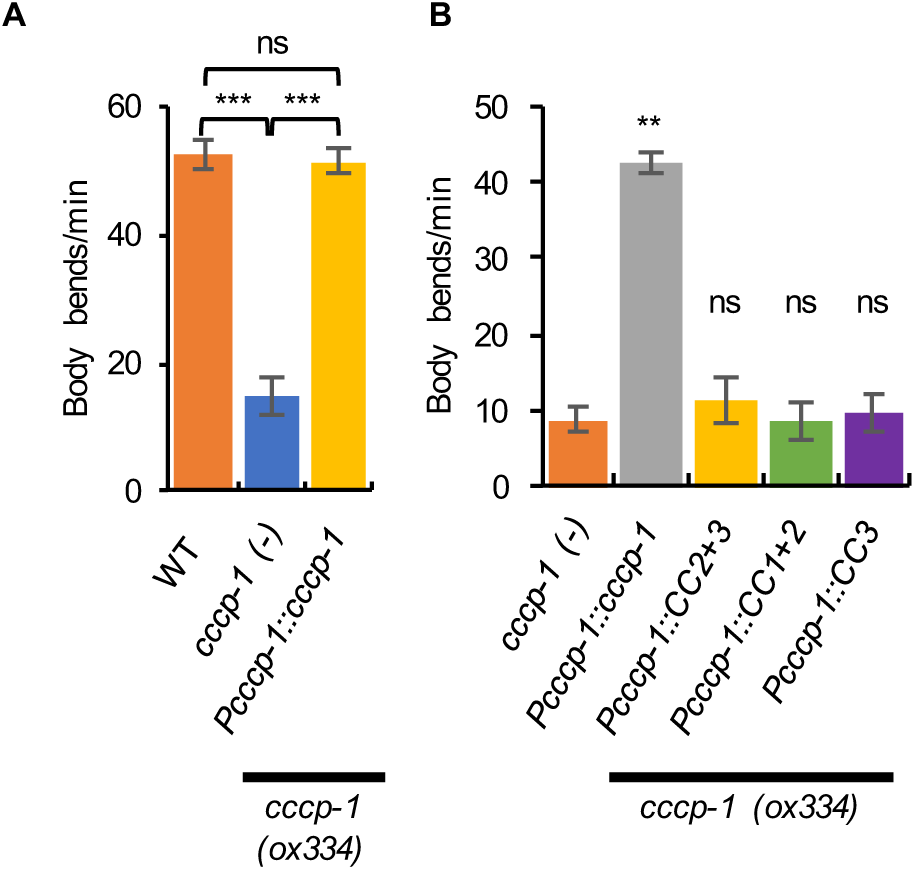
CC3 is necessary but not sufficient for CCCP-1 function in *C. elegans*. A, *cccp-1* mutant animals have slow locomotion which is rescued by a single-copy transgene of GFP-tagged CCCP-1 cDNA expressed under the *cccp-1* promoter. ***: p < 0.001, ns: p > 0.05, n=10 each. B, the slow locomotion of *cccp-1* mutants is not rescued by expression of CC3 or other CCCP-1 fragments. **: p < 0.01, ns: p > 0.05 compared to *cccp-1* mutant, n=10 each.

### 2.4 CCCP-1 binds membranes directly

The colocalization of CCCP-1 puncta with RAB-2 and trans-Golgi markers suggests that CCCP-1 may be localized to a membrane compartment, but CCCP-1 does not have a predicted transmembrane domain. To test whether CCCP-1 is a cytosolic protein associated with membranes, we used the rat insulinoma 832/13 cell line^33^. We transfected cells with a GFP-tagged rat CCCP1/CCDC186 construct and performed membrane fractionation assays (Figure 5A). Most of the CCCP1/CCDC186 protein associated with membranes (Figure 5A, P90 membrane pellet versus S90 soluble fraction). In the presence of either 1 M NaCl or 1% Triton X-100, CCCP1/CCDC186 was soluble. CCCP1/CCDC186 therefore behaves as a peripheral membrane protein.

**FIGURE 5.**
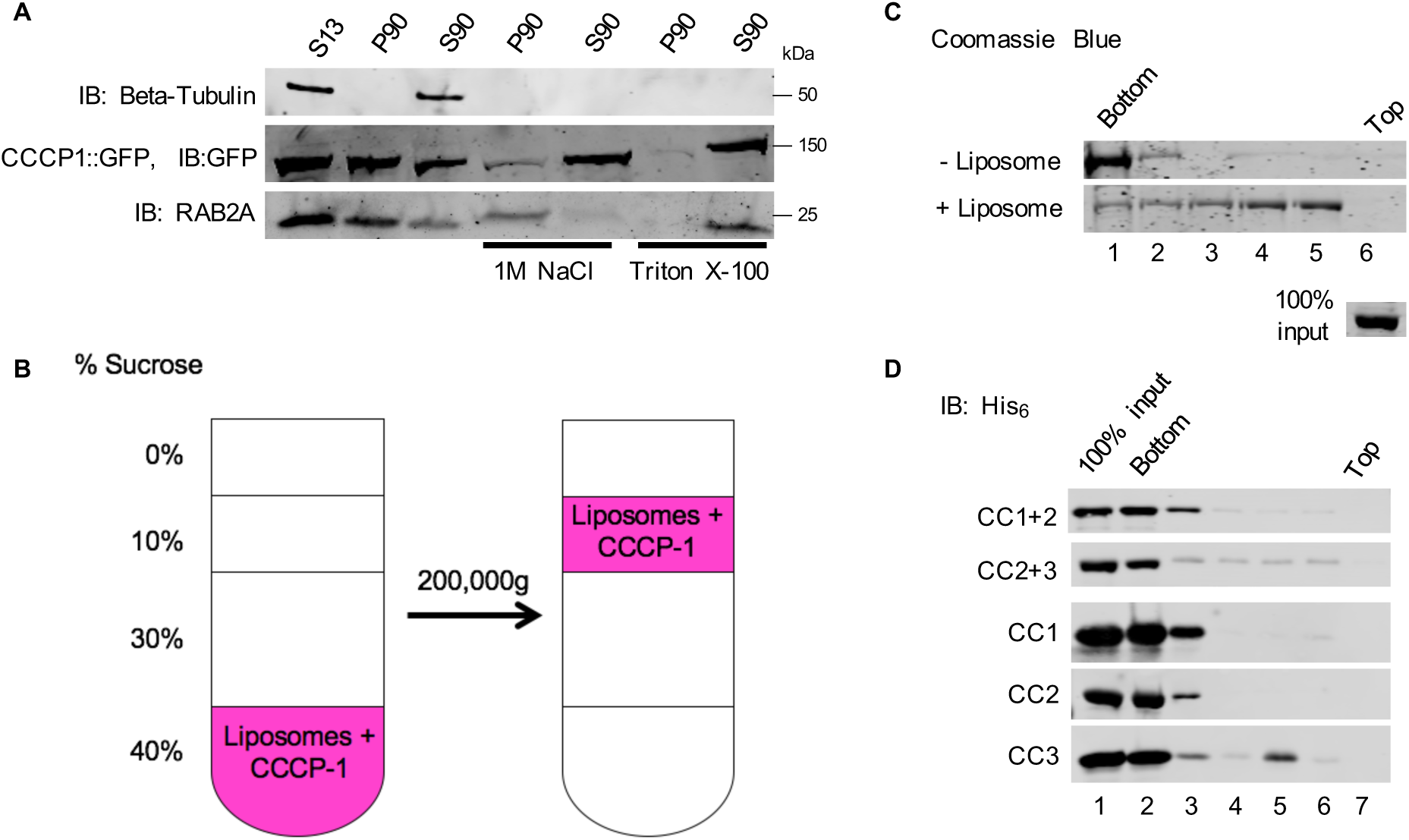
CC3 binds membranes directly. A, Rat CCCP1/CCDC186 is a peripheral membrane protein. In insulinoma 832/13 cell fractions, rat CCCP1/CCDC186 was found primarily in the post-nuclear P90 membrane fraction and could be extracted by 1 M NaCl or Triton X-100. RAB2A associates with membranes via a lipid anchor and could be extracted by Triton X-100 but not by 1 M NaCl. Tubulin served as a control soluble protein. S13: the supernatant obtained by a 13,000g spin of the post-nuclear supernatant, containing the cytosolic proteins and proteins associated with small membrane compartments. S90, P90: supernatant and pellet fractions obtained by a 90,000g spin of the S13 fraction, containing cytosolic and membrane-associated proteins respectively. B, Schematic of the liposome flotation assay. The CCCP-1 protein (1 µM) was incubated with Golgi-mix liposomes (1 mM lipids) containing a rhodamine-labeled lipid. The suspension was adjusted to 40% w/v sucrose and overlaid with three cushions of decreasing sucrose concentration. The tube was centrifuged for two hours at 200,000g. The efficiency of the flotation is demonstrated by observing the rhodamine-labeled lipid near the top of the tube. C, CCCP-1 binds to membranes directly. CCCP-1 membrane-binding activity was assayed by liposome flotation. Untagged recombinant CCCP-1 was assayed in the absence (top) or presence (bottom) of Golgi-mix liposomes. After centrifugation, fractions were collected from the top (lane 6) to the bottom (lane 1) and analyzed by Coomassie-stained SDS-PAGE. The protein input shown here was part of the same Coomassie gel. D, CC3 is necessary and sufficient for CCCP-1 membrane association. Representative blots from flotation assays using Golgi-mix liposomes and His_6_-tagged recombinant CCCP-1 fragments. After centrifugation, fractions were collected from the top (lane 7) to the bottom (lane 2) and analyzed by Western blot against the His_6_ tag.

Membrane association of CCCP-1 could occur via binding an additional protein or through its direct interaction with the phospholipid bilayer. We assayed whether the purified CCCP-1 protein binds lipids directly in blot overlay and liposome flotation experiments. We generated liposomes containing cholesterol and lipids with neutral and charged head groups that mimic the lipid composition of the Golgi^34^. The liposomes were doped with a fluorescent lipid, rhodamine-phosphatidylethanolamine (Rh-PE). We incubated liposomes and protein at the bottom of a stepped sucrose density gradient (Figure 5B). When centrifuged, most of the Rh-PE-labeled liposomes floated to near the top of the gradient, and most of the CCCP-1 protein migrated with the liposomes (Figure 5C, lanes 4 and 5). In the absence of liposomes, CCCP-1 remained at the bottom of the gradient. Thus, CCCP-1 binds membranes directly.

Domain structure prediction software does not indicate the presence of any canonical lipid-binding domain in CCCP-1, nor does CCCP-1 have a lipid-binding Amphipathic Lipid Packing Sensor (ALPS) motif (our unpublished observations). Thus, CCCP-1 may contain a previously uncharacterized lipid-binding domain or motif. To test whether CCCP-1 has affinity for phospholipids bearing specific head groups, we used a protein-lipid overlay assay (PIP strip) (Figure S1A). CCCP-1 showed a weak preference for binding to phosphatidylinositides with a single phosphate group: phosphatidylinositol 3-phosphate (PI(3)P), phosphatidylinositol 4-phosphate (PI(4)P) and phosphatidylinositol 5-phosphate (PI(5)P). In addition, CCCP-1 showed some affinity for phosphatidic acid (PA) and phosphatidylserine (PS) (lipid head groups with a negative net charge), and phosphatidylethanolamine (PE) (a neutral head group) (Figure S1A). Thus, at least in blot overlay assays, the interaction is not selective for a specific lipid and not purely electrostatic or hydrophobic. To determine whether CCCP-1 prefers binding to a specific phosphatidylinositol with a single phosphate, we performed a protein-lipid overlay assay using a membrane spotted with a range of concentrations of phosphatidylinositols (PIP array). No obvious chemical selectivity was apparent (Figure S1B).

Because CC3 is necessary and sufficient for CCCP-1 localization in vivo, we tested whether CCCP-1 binds membranes through CC3. In flotation assays, fragments containing CC3 (that is, CC3 or CC2 + 3) floated with the liposomes (Figure 5D, lanes 5 and 6). Other CCCP-1 fragments remained mostly at the bottom; however, a small amount of CC1 + 2 migrated to the top of the tube (Figure 5D, lanes 5 and 6). Because less CC3 bound to liposomes than full-length CCCP-1, the binding affinity of CC3 to membranes may be weaker. However, the CC3 protein was less stable than the full-length CCCP-1 so the decreased liposome binding of CC3 may also be due to its decreased stability. We conclude that CCCP-1 has the capacity to bind membrane bilayers via its CC3 domain. Together, these results demonstrate that CC3 contains a RAB-2 binding site and suggest the existence of a novel membrane-association domain or motif.

### 2.5 CCCP-1 forms oligomers and has an elongated structure

CCCP-1 is predicted to be a coiled-coil protein with a domain structure reminiscent of the golgins, elongated proteins that act as Golgi-localized tethering factors^28–32^. To test whether CCCP-1 has biophysical properties similar to golgins, we studied bacterially expressed *C. elegans* His_6_-tagged CCCP-1 (Figure 6A). Though CCCP-1 has a predicted monomeric molecular mass of 89 kDa, it eluted from a Superose 6 size-exclusion column in two peaks, both with an apparent mass larger than a 670 kDa globular protein standard (Figure 6B). This result suggests that CCCP-1 may have an elongated non-globular shape, form oligomers, or be aggregated. Golgins are elongated and generally form oligomers^28,29,35^. To study the biophysical properties of the protein further, we subjected His_6_-CCCP-1 to velocity sedimentation through a 5-25% linear sucrose gradient. His_6_-CCCP-1 sedimented in a broad peak with most of the protein migrating between the 44 kDa and 157 kDa standards (Figure 6C, lanes 3 and 4). Very little protein was found in the pellet (Figure 6C, lane 7), indicating that the protein is not aggregated. The combination of large apparent molecular weight by size-exclusion chromatography (SEC) and slow sedimentation suggests that CCCP-1 has an elongated shape. To test whether CCCP-1 exists in multiple populations of oligomeric states or conformations, we subjected SEC fractions 10 and 12 to velocity sedimentation in a sucrose gradient (Figure 6D). Fraction 10 sedimented faster than fraction 12 suggesting that the protein exists in multiple conformations or oligomeric states.

**FIGURE 6.**
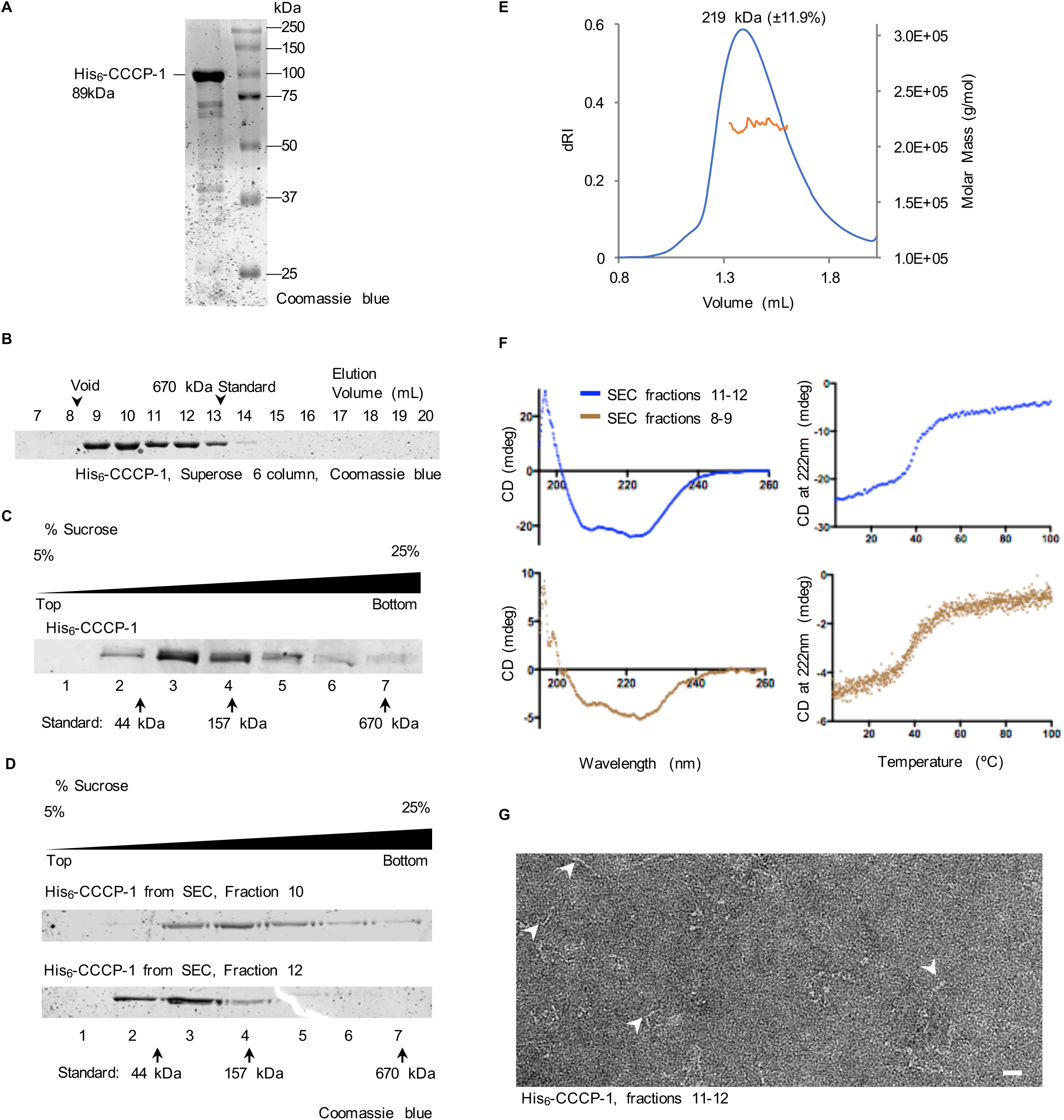
The CCCP-1 protein is elongated, alpha-helical and forms oligomers. A, Coomassie blue stained gel showing bacterially expressed His_6_-CCCP-1. B, recombinant His_6_-CCCP-1 protein runs larger by gel filtration than its predicted molecular weight. His_6_-CCCP-1 eluted in two peaks larger than the 670 kDa thyroglobulin standard. C, Velocity sedimentation using a 5-25% linear sucrose gradient shows His_6_-CCCP-1 sediments as a broad peak near its expected molecular weight, mainly between the 44 kDa ovalbumin and the 157 kDa gamma-globulin standards. After centrifugation, fractions were collected from top to bottom and analyzed by SDS-PAGE. D, the two peaks seen on SEC have different velocity sedimentation profiles suggesting a change in shape or a change in oligomeric state. Velocity sedimentation analysis of fractions 10 and 12 from size exclusion chromatography (see panel B) on a sucrose gradient identical to panel C. E, SEC-MALS analysis suggests that the smaller form of CCCP-1 is dimeric. The oligomeric state of His_6_-CCCP1 SEC fraction 12 was determined by SEC–MALS. The measured molar mass is 219 kDa (±11.9%). F, CCCP-1 is folded and composed mostly of alpha helices. Fractions eluting off the Superose 6 column (panel B) were analyzed by circular dichroism (CD) spectroscopy. (Left) Far-UV CD spectra of His_6_-CCCP-1. Ellipticity (mdeg) was plotted as a function of wavelength (nm). (Right) Melting curve spectra. Ellipticity (mdeg) was measured as a function of increasing temperature. G, CCCP-1 forms elongated and floppy filaments. Negative-stain electron microscopy of His_6_-CCCP-1. Protein eluting off a Superose 6 column (fractions 11 and 12) was imaged by electron microscopy. White arrowheads point to the elongated flexible structures. Scale bar: 20 nm.

To determine the oligomeric state, we analyzed His_6_-CCCP1 by SEC-Multiple Angle Light Scattering (SEC-MALS). Protein from SEC fraction 12 (Figure 6B) eluted as a monodisperse peak (polydispersity was 1 ± 16.8%) with a molar mass of 219 kDa (±11.9%) (Figure 6E). Thus, the smaller form of CCCP-1 is a dimer. Consistent with this, we found that CCCP-1 interacts with itself in yeast two-hybrid experiments (data not shown). We could not analyze the larger forms of the protein by SEC-MALS because they eluted in the column’s void.

To test whether the CCCP-1 protein contains mostly coiled-coils as predicted, we performed circular dichroism (CD) spectroscopy. The CD scans of the protein eluting in the pooled SEC fractions 11-12 exhibit the typical profile of a predominantly α-helical protein with minima at 208 nm and 222 nm (Figure 6F, top left), confirming that CCCP-1 is composed of the expected secondary structure. The melting curve shows a single sharp transition (Figure 6F, top right), implying that the protein is homogeneously folded. Moreover, the CD scans and melting curves of the protein eluting in the SEC void peak (fractions 8-9) are highly similar to those of the protein in fractions 11-12 (Figure 6F, bottom).

We further examined CCCP-1 structure by negative-stain electron microscopy (EM). CCCP-1 formed elongated filaments of varying shapes and sizes (Figure 6G), explaining why the protein runs large on SEC and sediments slowly. This result further suggests that CCCP-1 may exist as polymorphic oligomers with multiple flexible forms. Similar biophysical features have been observed for other golgins^35–37^.

To determine which domain of the protein is responsible for its large apparent molecular mass, we analyzed purified CCCP-1 fragments by SEC. Fragments containing the CC2 middle domain (CC2, CC1 + 2, and CC2 + 3) ran at a very large apparent molecular weight (Figure S2). Furthermore, we observed that CC1 and CC3 ran closer to but still larger than their predicted globular monomeric masses (Figure S2).

Together, these results indicate that CCCP-1 likely forms extended oligomers and that the CC2 middle domain mediates the formation of higher-order oligomeric states, the formation of elongated structures, or both.

### 2.6 CC3 is highly conserved among metazoans and beyond

In Basic Local Alignment Search Tool (BLAST) analyses, we found a single clear CCCP-1 ortholog in most metazoans, including primitive metazoans like sponges and cnidarians (Figure S3), though it is not apparent in other primitive metazoans including ctenophores and placozoans. Interestingly, we also found a likely CCCP-1 ortholog in two single-celled organisms closely related to metazoans: the choanoflagellate *M. brevicollis* and the snail symbiont *C. owczarzaki*. This suggests that CCCP-1 originated shortly before the origin of metazoans and was perhaps lost in some early metazoan lineages. An alignment of CCCP-1 and its orthologs shows that the C-terminal domain of CCCP-1 that includes the third coiled-coil domain and sequence downstream is the most highly conserved region of the protein (Figure S3). The middle part of the protein shows moderate conservation and the N-terminal region is the most diverged (Figure S3). Thus, the proposed lipid-binding and localization domain of CCCP-1 is the most highly conserved part of the protein, suggesting that it has been selected to maintain these functions and that the ancestral function of CCCP-1 may have been in membrane trafficking.

## 3. DISCUSSION

Here we performed a structure-function analysis of CCCP-1 to identify the important domains of the protein and assign activities to these domains. We found that the ∼200 amino acid C-terminal domain (CC3) has multiple activities. First, CC3 is necessary and sufficient for localization near immature DCVs and the trans-Golgi. Second, CC3 is necessary and sufficient for binding RAB-2. Third, CC3 mediates direct physical interaction with synthetic membrane bilayers. Furthermore, CC3 is required for CCCP-1 function in *C. elegans* neurons. We propose that CC3 is a novel membrane-binding domain that targets CCCP-1 to membranes and is necessary for CCCP-1 function in DCV biogenesis. Interestingly, RAB-2 is not required for the localization of either CCCP-1 or its CC3 domain in vivo. Thus, the CC3 domain appears to contain the information to target CCCP-1, even in the absence of RAB-2. Though CCCP-1 localization could theoretically be determined by its binding to membranes with specific physical or chemical properties, we found a relative lack of specificity in the interaction of CCCP-1 with lipids. Thus, CCCP-1 must also depend on additional factors for localization, perhaps through a combination of protein and lipid-binding activities.

### 3.1 CC3 directs CCCP-1 to immature DCVs near the trans-Golgi, potentially via a novel membrane-binding domain

Our data indicate that CC3 contains a membrane-binding domain and that direct membrane association may be important for the function of CCCP-1. Unlike other long coiled-coil trafficking proteins, CCCP-1 does not contain a recognizable lipid-binding domain such as the GRIP domain found in golgins (GRIP = “Golgi targeting domain golgin-97, RanBP2alpha, Imh1p and p230/golgin-245”)^24,28,29,38^. The blot overlay experiments suggest that CCCP-1 does not have chemical selectivity for a specific lipid and that it binds to both negatively-charged and neutral lipid head groups. These data raise the possibility that the interaction of CCCP-1 with membranes may depend on a combination of physical and chemical factors such as membrane curvature, electrostatics, hydrophobicity, or lipid packing. Protein-membrane interactions are often mediated by amphipathic helices such as the ALPS motif^39–41^. We found that CC3 contains several potential amphipathic helices in and downstream of its predicted coiled-coil domain (Figure S3, HeliQuest^42^), but none of them have the properties of the lipid-binding ALPS motif which contains bulky aromatic residues on the hydrophobic side of the helix and mostly noncharged residues on its polar side^34,40,43^.

### 3.2 Is CCCP-1 a golgin?

Several features of CCCP-1 suggest that it should be included as a member of the golgin family. First, CCCP-1 has a domain structure predicted to be largely coiled-coil, and golgins belong to a family of conserved proteins that are composed of extended coiled-coil domains^28,29^. Second, like other golgins, CCCP-1 is localized to the Golgi. Third, CCCP-1 localization is mediated by its C-terminal domain, in part through direct membrane association. Similarly, golgins are often anchored by their C-termini to specific regions of the Golgi^28,29^, either through a C-terminal transmembrane anchor or a lipid-binding domain^28,29^. Fourth, both CCCP-1 and golgins form oligomers^29,35,36,44^. Fifth, CCCP-1 forms elongated structures like golgins^29,35,36,44^. Sixth, both CCCP-1 and golgins bind activated Rab proteins.

What do these similarities suggest about the molecular function of CCCP-1? Golgins often function as molecular tethers, capturing incoming vesicles at the Golgi to help mediate fusion^28,29,45^, with different golgins tethering vesicles that emerge from different compartments^45^. Perhaps CCCP-1 also functions as a molecular tether to help facilitate membrane fusion events that occur during early steps in DCV biogenesis. Given that CCCP-1 is anchored at its C-terminus, interactions with other membrane compartments might be mediated by its N-terminal domains that may be extended away from the trans-Golgi.

### 3.3 What is the function of CCCP-1 in DCV biogenesis?

Why might a long coiled-coil tether be important in DCV biogenesis? There are several steps in DCV maturation that involve membrane fusion reactions that may require tethering molecules. One is the homotypic fusion of immature DCVs^4,18,46^, a process recently shown to involve the HID-1 protein^47^. Interestingly, *hid-1* mutants in *C. elegans* have defects in locomotion and DCV cargo sorting similar to *cccp-1* and *rab-2* mutants^21,48,49^, suggesting that RAB-2 and CCCP-1 may also be important for homotypic fusion of immature DCVs. A second step of DCV maturation that may involve membrane fusion reactions is the post-Golgi “sorting by exit” of non-DCV cargos from immature DCVs^2–4,50,51^. In *rab-2* mutants, DCV cargos are inappropriately lost to the endosomal/lysosomal system^19,20^, possibly due to overactivation of the sorting by exit pathway. Consistent with this possibility, we recently discovered that the endosomal-associated retrograde protein (EARP) complex functions in the RAB-2-CCCP-1 genetic pathway to mediate cargo sorting to DCVs^25^. EARP is a member of the family of multisubunit tethering complexes^25,52^. Perhaps CCCP-1 interacts with EARP to mediate the tethering of endosomally-derived vesicles that contain DCV cargo or recycled sorting factors.

## 4. MATERIALS AND METHODS

### 4.1 Strains

Worm strains were cultured and maintained using standard methods^53^. A complete list of strains and mutations used is provided in the Strain List (Table S1).

### 4.2 Molecular biology and transgenes

A complete list of constructs used in this study is provided in the Plasmid List (Table S2). Using the multisite Gateway system, we cloned the full-length *cccp-1b* cDNA tagged at the C-terminal with eGFP under the expression of the *rab-3* and *cccp-1* promoters into the pCFJ150 destination vector used for Mos1-mediated single copy insertion (MosSCI)^54,55^. For the different CCCP-1 truncations, vector backbones and *cccp-1* cDNA fragments containing 20-30 bp overlapping ends were PCR amplified and combined by Gibson cloning^56,57^. All single copy integrations were made by the direct injection MosSCI method at the *ttTi5605* insertion site^54,55^. Extrachromosomal arrays were made by standard transformation methods^58^. For most constructs, we isolated two or more independent lines that behaved similarly. We generated only one line for the constructs overexpressing CC3::GFP under the *rab-3* promoter, coexpressing tagRFP::RAB-2 and CCCP1::GFP, and coexpressing tagRFP::RAB-2 and CC3::GFP.

For bacterial protein expression, *C. elegans* cDNAs coding for RAB-2, CCCP-1, and CCCP-1 fragments (as shown in Figure 1B) were inserted in pGST or pHIS parallel vectors^59^ by Gibson cloning between the vector BamHI and XhoI restriction sites for RAB-2, and between the BamHI and EcoRI restriction sites for the other constructs.

For cloning the rat CCCP1 cDNA, rat cDNA was generated from PC12 cells using QuantiTect Reverse transcription kit (Qiagen) and an anchored Oligo(dT) primer. The CCCP1 cDNA was PCR amplified using gene specific primers and sequenced. The sequence had a few variations compared to the reported sequence and was submitted to Genbank (accession # KX954625). Rat CCCP1 and its fragments CC1 + 2 (amino acids 1-743) and CC3 (amino acids 750-922) were cloned into the pEGFP-N1 vector at the EcoRI and BamH1 restriction sites using either classical restriction digest and ligation for the full length protein or Gibson cloning with PCR amplified vector and inserts for the fragments.

### 4.3 *C. elegans* fluorescence imaging

Worms were mounted on 2% agarose pads and anesthetized with 100 mM sodium azide. To image the dorsal nerve cords, young adult animals were oriented with dorsal side up by exposure to the anesthetic for ten minutes before placing the cover slip. Images were obtained using a Nikon Eclipse 80i wide-field compound microscope with 40× or 60× oil objectives (numerical apertures of 1.30 and 1.40 respectively). Images were acquired at room temperature using an Andor Technology Neo sCMOS camera, model number DC152Q-C00-FI. The acquisition software used was NIS Elements AR 4.10.01. Raw images were then cropped with FIJI. Strains were imaged multiple times.

### 4.4 Cell culture and immunostaining of 832/13 cells

The insulinoma INS-1-derived 832/13 rat cell line was obtained from Dr. Christopher Newgard (Duke University School of Medicine) via Dr. Ian Sweet and Dr. Duk-Su Koh (University of Washington). 832/13 cells were routinely grown at 5% CO2 at 37°C in RPMI-1640, GlutaMAXTM (GIBCO), supplemented with 10% FBS, 1 mM Sodium pyruvate, 10 mM HEPES, 50 µM 2-mercaptoethanol, and 1X Pen/Strep (GIBCO). Cells were passaged biweekly after trypsin-EDTA detachment. All studies were performed on 832/13 passages between 70 and 90.

832/13 cells were plated on a coverslip at 80–90% confluency. After 24 to 48h hours, cells were transfected with a construct expressing C-terminally EGFP-tagged rat CCCP1, CC1 + 2 (amino acids 1-743) or CC3 (amino acids 750-922) using Lipofectamine 2000 (Thermo Fisher) according to the manufacturer’s instructions. Cells were immunostained as described^25^. Primary antibodies were the mouse monoclonal anti-GM130 (1:200, BD biosciences, #610822), mouse monoclonal anti-Chromogranin A (1:50, Santa Cruz, #sc-393941), rabbit polyclonal anti-TGN38 (1:350, Sigma #T9826) rabbit polyclonal anti-GFP (1:200, Santa Cruz, #sc-8334), mouse monoclonal anti-GFP (1:200, Santa Cruz #sc-9996). Secondary antibodies were the Rhodamine anti-rabbit secondary antibody (1:1000, Jackson Immunoresearch #111-025-144) and Alexa Fluor 488 anti-mouse secondary antibody (1:1000, Jackson Immunoresearch #115-545-146). Images were obtained using an Olympus FLUOVIEW FV1200 confocal microscope with a 60× UPlanSApo oil objective with a numerical aperture of 1.35. The acquisition software used was Fluoview v4.2. Pearson’s correlation coefficients were determined using FIJI and the JaCOP plugin by drawing a rectangle around the perinuclear structure for each cell.

### 4.5 Locomotion assays

To measure worm locomotion, first-day adults were picked to thin lawns of OP50 bacteria (2-3 day-old plates) and body bends were counted for one minute immediately after picking. A body bend was defined as the movement of the worm from maximum to minimum amplitude of the sine wave. Worms mutant for *cccp-1*, like other mutants affecting DCV function, have a stereotypical “unmotivated” phenotype in which worms are slow when placed on food^24^. In our assay conditions in which the worm was stimulated by transfer to a new plate, expression of full length CCCP-1 fully rescued the locomotion defect of *cccp-1* mutants (Figure 4A). However, we observed that the transgene only partially rescued the locomotion defect of a *cccp-1* mutant when worms were not stimulated. This incomplete rescue could be due to the GFP tag, differences in expression levels, or the use of cDNA in the transgenes. This result further suggests that worm locomotion is sensitive to CCCP-1 expression levels. The locomotion assays were repeated twice or more.

### 4.6 Protein expression in bacteria and purification

GST-RAB-2, His_6_-CCCP-1, GST-CCCP-1, GST, and all His_6_-CCCP-1 fragments were transformed into *E. coli* BL21 (DE3), pLys Rosetta cells (Invitrogen). His_6_-CCCP-1 was grown in LB medium containing ampicillin and chloramphenicol to OD600 = 0.4−0.6. Protein expression was induced with 0.4 mM Isopropyl β-D-1-thiogalactopyranoside (IPTG) at 20°C overnight. Bacteria were harvested by a 5,000g spin at 4°C and cells were resuspended in ice-cold lysis buffer containing 50 mM Tris, pH 7.6, 200 mM NaCl, 10% glycerol, 5 mM 2-mercaptoethanol, 1-2 mM MgCl_2_, 0.5-1% Triton-X100, up to 1 µL of benzonase nuclease (Sigma) per 10 mL of lysis buffer, and supplemented with protease inhibitor cocktail (Pierce, according to manufacturer’s instructions) and 1 mM PMSF. 10-25 mM imidazole was added to the lysis buffer to decrease nonspecific binding to the nickel resin. Cells were lysed by sonication. An additional 1 mM PMSF was added to the lysate during sonication. Lysates were clarified by a 20,000g spin at 4°C. The supernatant was incubated with PerfectPro Ni-NTA Agarose (5 Prime) by either gravity flow using a disposable column or batch purification. The resin was washed with buffer containing 50 mM Tris, pH 7.6, 25-35 mM imidazole, 200 mM NaCl, 10% glycerol, and 5 mM 2-mercaptoethanol. The resin was eluted with the same buffer containing 250 to 400 mM imidazole.

For the His_6_-CCCP-1 used for SEC, CD and EM, the protein eluted from the Nickel-NTA beads was concentrated using an Amicon-Ultra centrifugal filter, and buffer exchanged to 50 mM Tris pH 7.6, 200 mM NaCl, 5 mM 2-mercaptoethanol using a PD-10 column (GE Healthcare). Protein aliquots were flash-frozen in liquid nitrogen and stored at −80°C.

The His_6_ tag of full length CCCP-1 was cleaved off using TEV protease, which was expressed and purified as described^60^. An estimated 1:10 molar amount of His_6_-TEV protease was added to the eluted CCCP-1 protein and incubated overnight at 4°C with gentle rotation. The protein was buffer exchanged (20 mM Tris pH 7.6, 200 mM NaCl, 2 mM 2-mercaptoethanol) by dialysis. The cleaved tag and the protease were removed by incubation with PerfectPro Ni-NTA resin for 1 hour at 4°C. Cleaved protein was concentrated, supplemented with 10% glycerol, aliquoted and flash frozen in liquid nitrogen.

His_6_-CCCP-1 fragments were grown in TB medium containing ampicillin and chloramphenicol to OD600 = 0.8-1 and protein expression was induced with 1 mM IPTG. The protein was handled like His_6_-CCCP-1 full length except that the lysis buffer lacked benzonase and MgCl_2_. The eluted protein was dialyzed with 20 mM Tris pH 7.6, 200 mM NaCl, 2 mM 2-mercaptoethanol, concentrated, supplemented with 10% glycerol, aliquoted and flash frozen in liquid nitrogen.

For the GST-CCCP-1 and GST used for the protein-lipid overlay assays, cells were grown in TB medium containing ampicillin and chloramphenicol to OD600 = 0.5-0.8 and protein expression was induced with 0.5 mM IPTG. After induction, bacteria were then incubated at 20°C overnight. Harvested cells were resuspended in the same ice-cold lysis buffer described above for His_6_-tagged proteins except lacking imidazole, benzonase and MgCl_2_ and lysed by sonication. The clarified lysate was incubated with GST-Sepharose resin (GE Healthcare). The resin was then washed with 50 mM Tris, pH 7.6, 200 mM NaCl, 10% glycerol, 5 mM 2-mercaptoethanol and eluted with the same buffer containing 20 mM reduced glutathione. The eluted protein was dialyzed (20 mM Tris pH 7.6, 200 mM NaCl, 2 mM 2-mercaptoethanol, supplemented with 10% glycerol), aliquoted and flash frozen in liquid nitrogen.

GST-RAB-2 and GST used for the GST-RAB-2 pull downs were grown in LB medium containing ampicillin and chloramphenicol to OD600 = 0.4-0.6. Protein expression was induced with 1 mM IPTG at 18°C overnight. Bacteria were harvested and resuspended in ice-cold lysis buffer containing 50 mM Tris, pH 7.6, 200 mM NaCl, 10% glycerol, 5 mM 2-mercaptoethanol, 2 mM MgCl_2_, 0.2% Triton-X100, up to 1 µL of benzonase nuclease (Sigma) per 10 mL of lysis buffer and supplemented with protease inhibitor cocktail (Pierce, according to manufacturer’s instructions) and 1 mM PMSF. Cells were lysed as above and stored as a clarified lysate at −80°C. The concentration of GST or GST-RAB-2 in the lysate was estimated by a small-scale affinity purification.

Protein expression and purification conditions typically yielded more than 1 mg of protein per liter of culture media. The protein concentration was measured with Bradford reagent and purity was assessed by Coomassie-stained SDS-PAGE. GST-tagged proteins were estimated to be over 95% pure. CCCP-1 fragments containing CC3, especially CC3 alone, yielded less protein and the measured concentration of protein was adjusted by comparing the band intensity with purer His_6_-tagged fragments on a Western blot. All proteins migrated on SDS-PAGE to their expected molecular weight, except for GST-RAB-2 (predicted 50 kDa) that migrated between the 37 kDa marker and the 50 kDa marker, and the His_6_-CC1 fragment (predicated 19 kDa) that migrated above the 20 kDa marker.

### 4.7 Western blotting

Protein samples were solubilized in SDS loading dye and resolved on 8%, 10% or 12% SDS-PAGE gels. Proteins were then transferred to PVDF or nitrocellulose membranes using a semi-dry transfer apparatus (Biorad) system. The membranes were blocked with 3% dry milk in TBST (10 mM Tris pH 7.4, 150 mM NaCl, 0.05% Tween-20) for 1 hour at room temperature or overnight at 4°C. Primary and secondary antibodies were diluted in TBST + 3% dry milk and incubated 1 hour at room temperature or overnight at 4°C. The following primary antibodies were used: mouse monoclonal anti-His_6_ (1:1000, Thermo Scientific HIS.H8 #MA1-21315), mouse monoclonal anti-GFP (1:1000, Roche #11814460001), mouse monoclonal anti-beta-tubulin antibody (1:1000, Thermo Scientific #MA5-16308) and rabbit polyclonal anti-Rab2 (1:200, Santa Cruz Biotechnology (FL-212) sc-28567, produced from a human RAB2A antigen). The secondary antibodies were: Alexa Fluor 680-conjugated affinity pure goat anti-mouse antibody (1:20,000, Jackson Immunoresearch #115-625-166) and Alexa Fluor 680-conjugated affinity pure goat anti-rabbit antibody (1:20,000 Jackson Immunoresearch #111-625-144). A Li-COR processor was used to develop images.

### 4.8 GST-RAB-2 pulldowns

To decrease nonspecific binding, we used LoBind microcentrifuge tubes (Eppendorf). 25 µL of glutathione sepharose resin (GE Healthcare) was blocked for 1 hour at room temperature or overnight at 4°C with 5% bovine serum albumin (BSA) in reaction buffer (50 mM Tris pH 7.6, 150 mM NaCl, 10% glycerol, 5 mM 2-mercaptoethanol, 5 mM MgCl_2_ and 1% Triton-X100). The resin was washed with reaction buffer and incubated with clarified bacterial lysates containing about 50 µg of GST-RAB-2 or 25 µg of GST for 1 hour at 4°C with gentle agitation. Beads were washed with nucleotide loading buffer (50 mM Tris pH 7.6, 150 mM NaCl, 10% glycerol, 5 mM 2-mercaptoethanol, 5 mM EDTA and 1% Triton-X100) and incubated with either 60X molar ratio of GTPγS or 300X of GDP on a rotator at room temperature for 2 hours. 20 mM MgCl_2_ was added to the reaction and incubated for another 15 minutes at room temperature. Nucleotides were washed off with reaction buffer and the beads were incubated with CCCP-1 or its fragments (at 1:2 molar ratio with GST-RAB-2) for 2 hours at 4°C on a rotator. Beads were washed three times with low volumes of reaction buffer (3 × 200 µL) and the resin was eluted with 20 mM reduced glutathione in reaction buffer. Half of the eluent was loaded on SDS-PAGE gels and analyzed by Western blotting against the His_6_-tag. The pulldown was repeated four times with the full-length CCCP-1 protein and twice with the fragments with identical results.

### 4.9 Cell fractionation

We used a similar method to the one described^52^. Specifically, 832/13 cells were grown on a 15 cm dish to 90% confluence and transfected with CCCP1::eGFP using Lipofectamine 2000 (Thermo Fisher) according to the manufacturer’s instructions. After 24 h, cells were washed twice with ice cold PBS and detached using a cell scraper. Cells were then transferred to a microcentrifuge tube and pelleted for 5 min at 300g in a tabletop centrifuge at 4°C. Cells were resuspended in 500 µL sucrose buffer (20 mM HEPES, pH 7.4, 250 mM sucrose supplemented with protease inhibitors (Pierce) and 1 mM PMSF) and were homogenized on ice by a Dounce homogenizer (20 strokes). The cell lysate was then centrifuged at 1,000g for 5 minutes at 4°C to remove unbroken cells and nuclear debris. The post-nuclear supernatant was further clarified by centrifugation at 13,000g for 10 min at 4°C. The supernatant was divided into four samples, one of which was supplemented with SDS loading buffer (supernatant, S13). Membranes in the other three samples were pelleted using a Beckman TLA100 rotor (45 minutes, 90,000g, 4°C). The supernatant from one of the samples was supplemented with SDS page loading buffer (supernatant, S90) and its pellet was resuspended in an equal volume of sucrose buffer and supplemented with SDS loading buffer (pellet, P90). The pellets of the two last samples were resuspended in either high-salt buffer (50 mM Tris, pH 7.4, 1 M NaCl, 1 mM EDTA supplemented with 10% glycerol and protease inhibitors) or detergent buffer (50 mM Tris, pH 7.4, 150 mM NaCl, 1% Triton X-100, 1 mM EDTA supplemented with 10% glycerol and protease inhibitors). After incubation for 1 hour on ice the samples were centrifuged at 90,000g for 45 min at 4°C. The collected membrane fractions, either salt-extracted (S90, 1 M NaCl) or detergent-extracted (S90, 1% Triton X-100), were then supplemented with SDS loading buffer. Their pellets were resuspended in equal volumes of high salt or detergent buffer and then supplemented with SDS loading buffer (P90, 1 M NaCl and P90, 1% Triton X-100). Samples were analyzed by Western blot.

### 4.10 Golgi-mix liposome preparation

The following lipids were used: 1-palmitoyl-2-oleoyl-sn-glycero-3-phosphocholine (POPC), 1,2-dipalmitoyl-sn-glycero-3-phosphoethanolamine (DPPE), and 1,2-di-oleoyl-sn-glycero-3-phospho-l-serine (DOPS) were purchased from NOF America Corporation. Cholesterol, L-α-phosphatidylinositol from soy (PI) and 1,2-dioleoyl-sn-glycero-3-phosphoethanolamine-N-(lissamine rhodamine B sulfonyl) (Rh-PE) were purchased from Avanti Polar Lipids. The composition of Golgi-mix liposomes was as described^34^ with the following fractions given as molar percent: POPC (49%), DPPE (19%), DOPS (5%), Soy-PI (10%), cholesterol (16%) and Rhodamine-PE (1%). The lipids dissolved in chloroform were mixed together and chloroform was dried under compressed nitrogen for 3 hours to overnight and then lyophilized in a Labconco Free Zone 2.5 lyophilizer for 1 hour to remove residual chloroform. The dried lipid mixture was resuspended in 50 mM Tris pH 7.6 and 50 mM NaCl. To generate the liposomes, the mixture was sonicated for 5 minutes in a 50°C water bath. Liposomes were stored at room temperature and used within 2-3 days.

### 4.11 Flotation experiments

Proteins (1 µM) and Golgi-mix liposomes (1 mM) were incubated in 50 mM Tris pH 7.6, 50 mM NaCl buffer at room temperature for 30 minutes. The suspension was adjusted to 40% sucrose by adding 75% w/v sucrose solution in the same buffer for a total volume of 200 µL and transferred to a polycarbonate ultracentrifuge tube (Beckman Coulter, 343778). Three layers were overlaid on top of the high sucrose suspension: 500 µL of 30% sucrose, 300 µL of 10% sucrose and 200 µL of 0% sucrose in 50 mM Tris pH 7.6, 50 mM NaCl. The sample was centrifuged at 200,000g in a Beckman Coulter swinging bucket rotor (TLS 55) for 2 hours at 25°C. Most of the fluorescently labeled liposomes migrated to the top of the tube in three layers close to the 0%-10% sucrose interface. Six fractions of 200 µL each were collected from the top. The second fraction contained most of the pink color coming from rhodamine, with some in the third fraction as well. Experiments with full-length untagged CCCP-1 were analyzed by SDS-PAGE and Coomassie blue staining. Experiments with the His_6_-tagged CCCP-1 fragments were analyzed by Western blot using an antibody to the His_6_ epitope tag. Experiments with the full-length CCCP-1 protein were performed three times with two independent liposome preparations. The key experiments with CC3 and CC1 + 2 were performed side by side twice with independent liposome preparations and both found CC3 to be a much stronger liposome binder.

### 4.12 PIP Strips and PIP arrays

The PIP strips and PIP arrays were purchased from Echelon and used as recommended by the manufacturer. The membranes were blocked for 1 hour at room temperature with PBST (0.1% v/v Tween-20) + 3% fatty acid-free BSA (Sigma). The membranes were incubated for 1 hour at room temperature with GST or GST-CCCP-1. For the PIP strip, equimolar amounts of GST and GST-CCCP-1 were used (around 100 nM in PBST + 3% fatty acid-free BSA). For the PIP array, 70 nM of GST-CCCP-1 and 150 nM of GST were used. The membranes were washed with PBST and incubated with the anti-GST antibody Horseradish Peroxidase (HRP) conjugate (K-SEC2 from Echelon) in PBST + 3% fatty acid-free BSA. Lipid binding was detected by adding 3,3’,5,5’-tetramethylbenzidine (TMB) precipitating solution (K-TMB from Echelon). The positive control PI(4,5)P2 Grip (PLC-δ1-PH) was obtained from Echelon (Catalog #G-4501) and used as recommended by the manufacturer. The membranes were imaged with a conventional digital camera. The experiments were performed once each.

### 4.13 Size-exclusion chromatography (SEC)

Aliquots of His_6_-CCCP-1 expressed and purified as described above were loaded on a Superose 6 column (GE Healthcare) at 4°C. His_6_-CCCP-1 was eluted with 50 mM Tris pH 7.6, 200 mM NaCl and 5 mM 2-mercaptoethanol. Given that CCCP-1 does not contain any tryptophan residues, its extinction coefficient is very low. Since the size-exclusion chromatogram was very sensitive to more optically absorbent contaminants, we collected 1 mL fractions and analyzed them by Coomassie-stained SDS-PAGE gels. Recombinant His_6_-CCCP-1 fragments were loaded on a Superose 6 (or Superdex 200) column and eluted with 50 mM Tris pH 7.6 and 200 mM NaCl. Fractions were analyzed by Western blot to the His_6_ tag.

### 4.14 Velocity Sedimentation

5-25% linear sucrose gradients were made in 50 mM Tris pH 7.6, 200 mM NaCl, 5 mM 2-mercaptoethanol in polycarbonate ultracentrifuge tubes (Beckman Coulter, 349622). Aliquots of His_6_-CCCP-1 SEC fractions 10 or 12 were loaded at the top of the gradient and tubes were centrifuged at 100,000g for 16 hours at 4°C in a Beckman Coulter swinging bucket rotor (SW 50). Fractions were collected from top to bottom and analyzed by SDS-PAGE and Coomassie blue staining.

### 4.15 SEC–MALS

SEC–MALS was performed by injecting fraction 12 from SEC (Figure 6B) of purified recombinant His_6_-CCCP-1 on a Superose 6 Increase 3.2/100 column (GE Healthcare) that was equilibrated in 50 mM Tris pH 7.6, 200 mM NaCl, 5 mM 2-mercaptoethanol. The eluted sample was monitored by ultraviolet absorption at 280 nm, light scattering at 658 nm (Wyatt Technology) and differential refractometry (Wyatt Technology). The data was analyzed using ASTRA software (Wyatt Technology). The protein absolute molecular mass was calculated assuming a dn/dc value of 0.185.

### 4.16 Circular dichroism spectroscopy

Samples were analyzed using a Jasco J-1500 circular dichroism (CD) spectrometer. Protein in 50 mM Tris pH 7.6, 200 mM NaCl and 5 mM 2-mercaptoethanol was used to collect the CD data. Fractions from SEC (pooled fractions 8-9 and 11-12, Figure 6B) were concentrated around 10-fold using an Amicon Ultra centrifugal unit and flash frozen in liquid nitrogen before use. The signal was blanked with buffer only. For the CD scan, the samples were kept at 4°C and the ellipticity (in mdeg) was measured at wavelengths between 195 nm and 260 nm. For the melting curve, the ellipticity (mdeg) was measured at 222 nm as the temperature was increased from 4°C to 100°C.

### 4.17 Negative-stain electron microscopy

Frozen aliquots of pooled and concentrated SEC fractions 11-12 were diluted to 0.01 mg/mL in 50 mM Tris pH 7.6, 200 mM NaCl and 5 mM 2-mercaptoethanol. Samples were prepared by negative stain, using a 0.7% uranyl formate solution, and spotted onto a 400-mesh copper-coated grid. Electron micrographs were acquired on a Tecnai T12 microscope (FEI co.), operating at 120 kV, at 52,000X magnification, with images taken by a Gatan US4000 CCD camera. Similar results were observed using protein from an independent protein preparation and imaged using a Morgagni M268 electron microscope.

### 4.18 Statistics

For the *C. elegans* locomotion assay, data were tested for normality by a Shapiro-Wilk test. We used the Kruskal-Wallis test followed by Dunn’s test to investigate whether there was statistical significance between groups. For the 832/13 cells, data were tested for normality by a Shapiro-Wilk test followed by a one-way Anova test with Bonferroni correction to investigate whether there was statistical significance between groups.

## Acknowledgments

We thank Kumud Raj Poudel and Jihong Bai for sharing purified phospholipids and for help with generating liposomes; Amanda Clouser for help with CD spectroscopy; Amanda Clouser and Mengtong Duan for help with SEC-MALS; Christopher Newgard, Ian Sweet, and Duk-Su Koh for the 832/13 and PC12 cell lines; Suzanne Hoppins for sharing equipment and experimental guidance; Maximillian Greenwald for assistance with *C. elegans* locomotion assays; Andrew Borst for help with electron microscopy; and Jihong Bai for comments on the manuscript. JC was supported in part by an NIH Institutional Training Grant for Neurobiology (T32 GM007108). This work was supported by a University of Washington Diabetes Research Center Pilot and Feasibility Award to MA (NIH grant P30 DK017047), by NIH grant R01 GM077349 to AJM, and by NIH grant R00 MH082109 and an Ellison Medical Foundation New Scholar Award to MA.

## Supplementary Figure Legends

**FIGURE S1.**
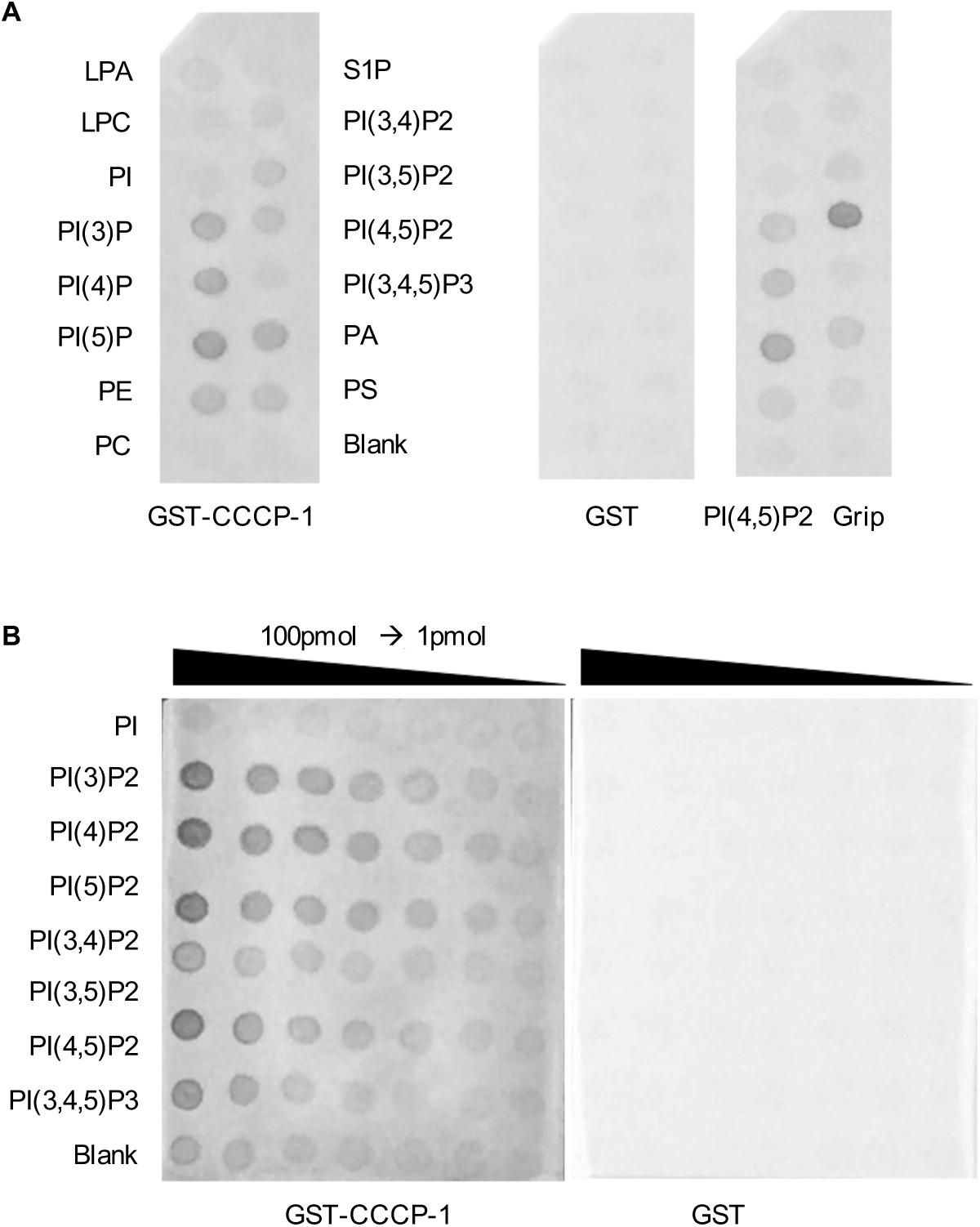
CCCP-1 binds to phosphatidylinositol lipids with a single phosphate group. A, Equimolar amounts of GST-CCCP-1 or GST were incubated with membranes coated with different membrane phospholipids (PIP strips). Binding activity was detected using an antibody to the GST tag. PI(4,5)P2 Grip, a GST-tagged PLC-δ1-PH domain protein, was used as a positive control. LPA: lysophosphatidic acid, LPC: lysophosphocholine, PI: phosphatidylinositol, PI(3)P: phosphatidylinositol (3) phosphate, PI(4)P: phosphatidylinositol (4) phosphate, PI(5)P: phosphatidylinositol (5) phosphate, PE: phosphatidylethanolamine, PC: phosphatidylcholine, S1P: sphingosine 1-phosphate, PI(3,4)P2: phosphatidylinositol (3,4) bisphosphate, PI(3,5)P2: phosphatidylinositol (3,5) bisphosphate, PI(4,5)P2: phosphatidylinositol (4,5) bisphosphate, PI(3,4,5)P3: phosphatidylinositol (3,4,5) trisphosphate, PA: phosphatidic acid, and PS: phosphatidylserine. B, CCCP-1 does not show obvious binding selectivity between phosphatidylinositol lipids. Equimolar amounts of GST-CCCP-1 or GST were incubated with membranes coated with different phosphatidylinositols of decreasing concentration (PIP arrays). Binding activity was detected using an antibody to the GST tag.

**FIGURE S2.**
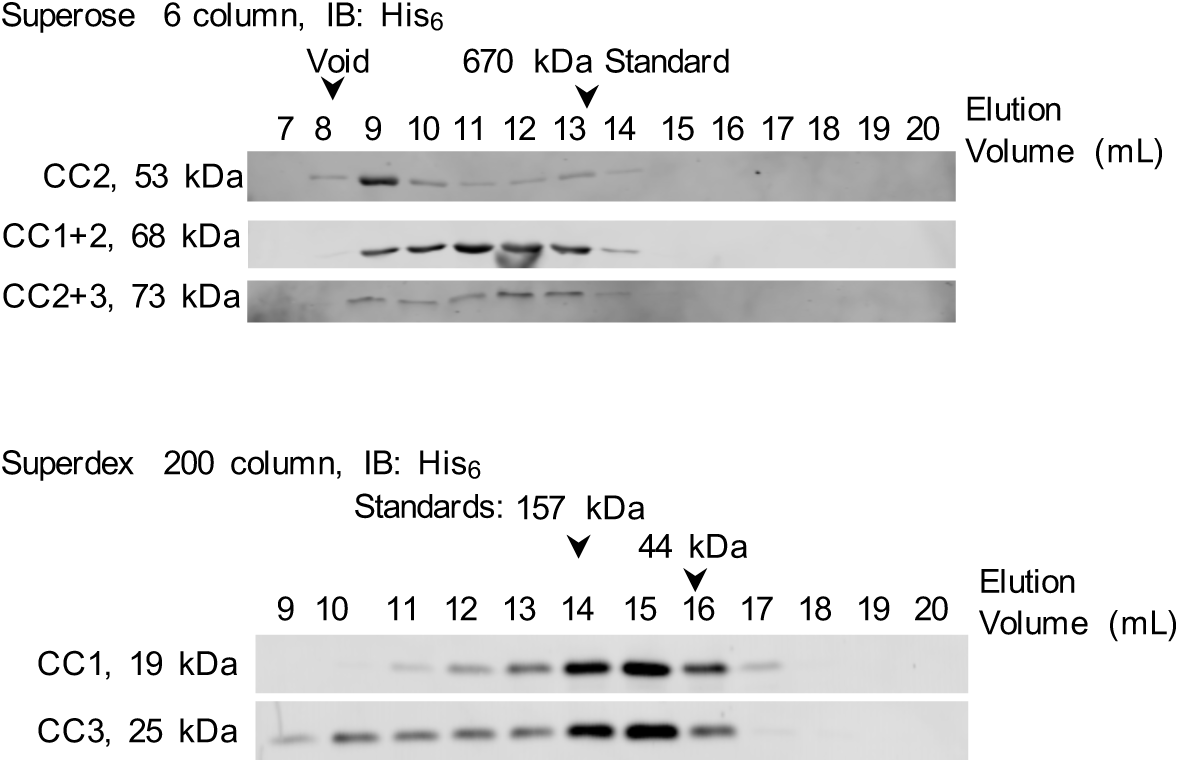
The central CC2 domain is responsible for the large apparent molecular mass of CCCP-1. Fractionation of CCCP-1 fragments by gel filtration on a Superose 6 column for large fragments (top) or Superdex 200 column for smaller fragments (bottom). 1 mL fractions were collected and analyzed by Western blot against the His_6_ tag. IB: immunoblot.

**FIGURE S3.**
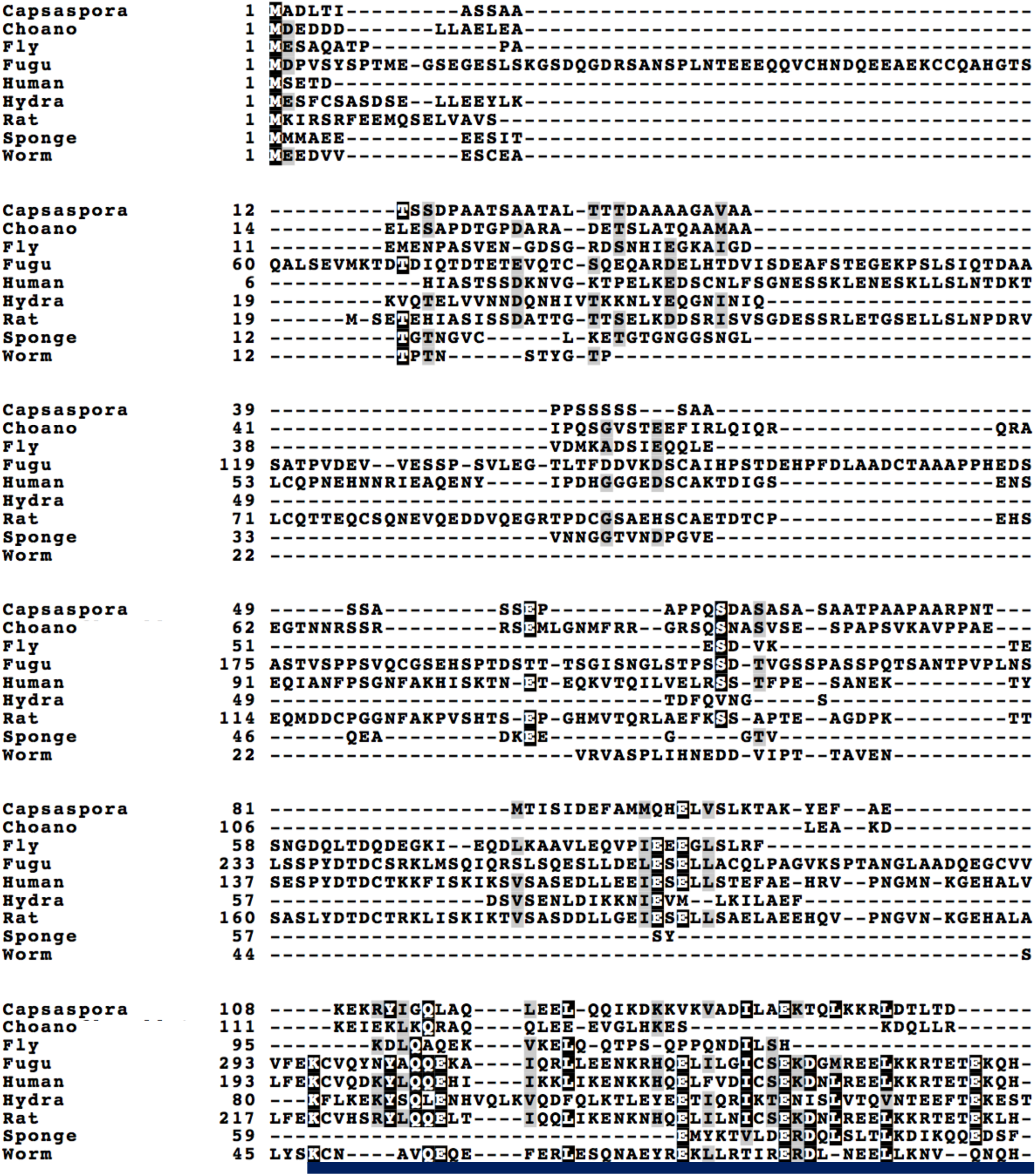
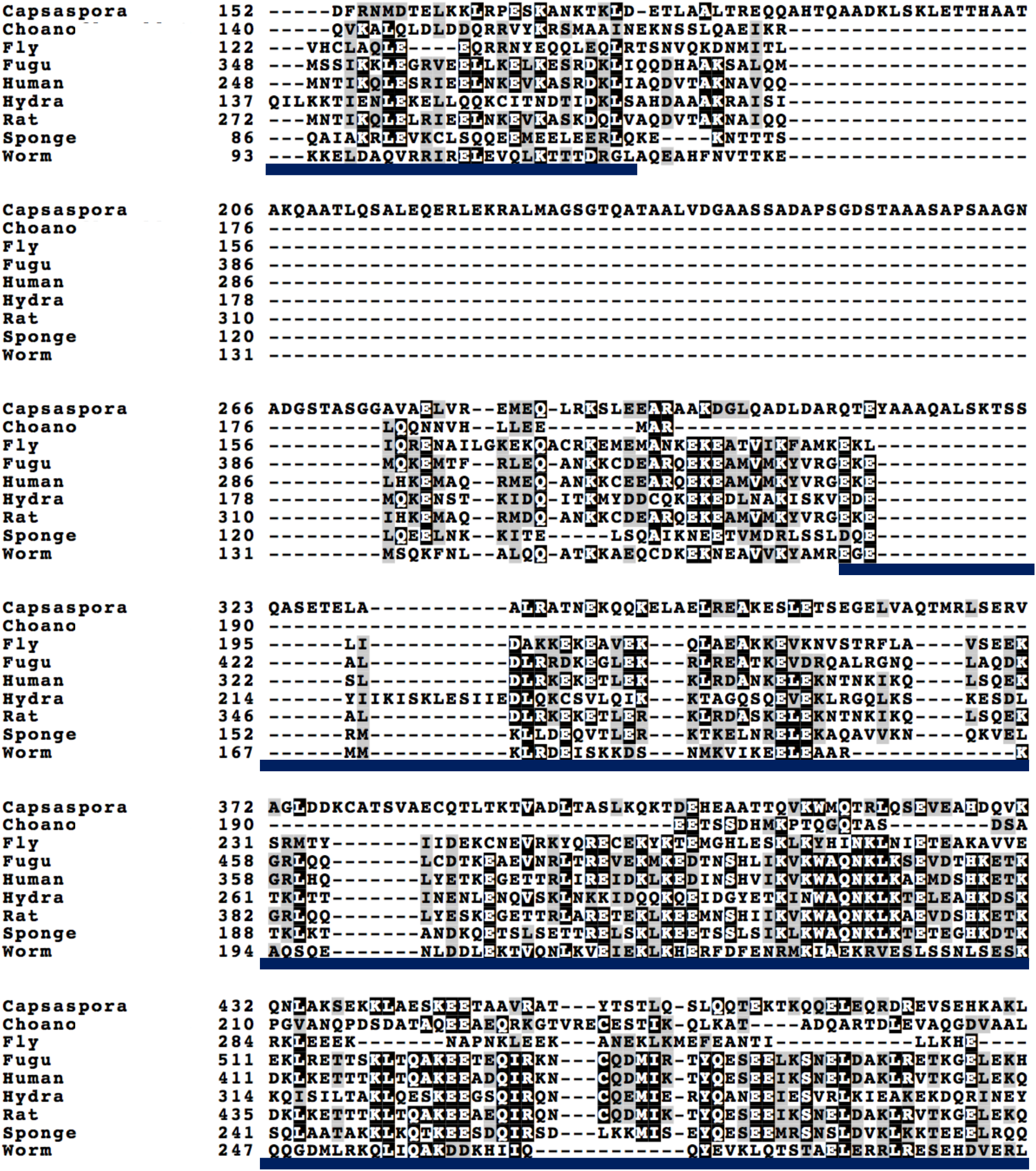
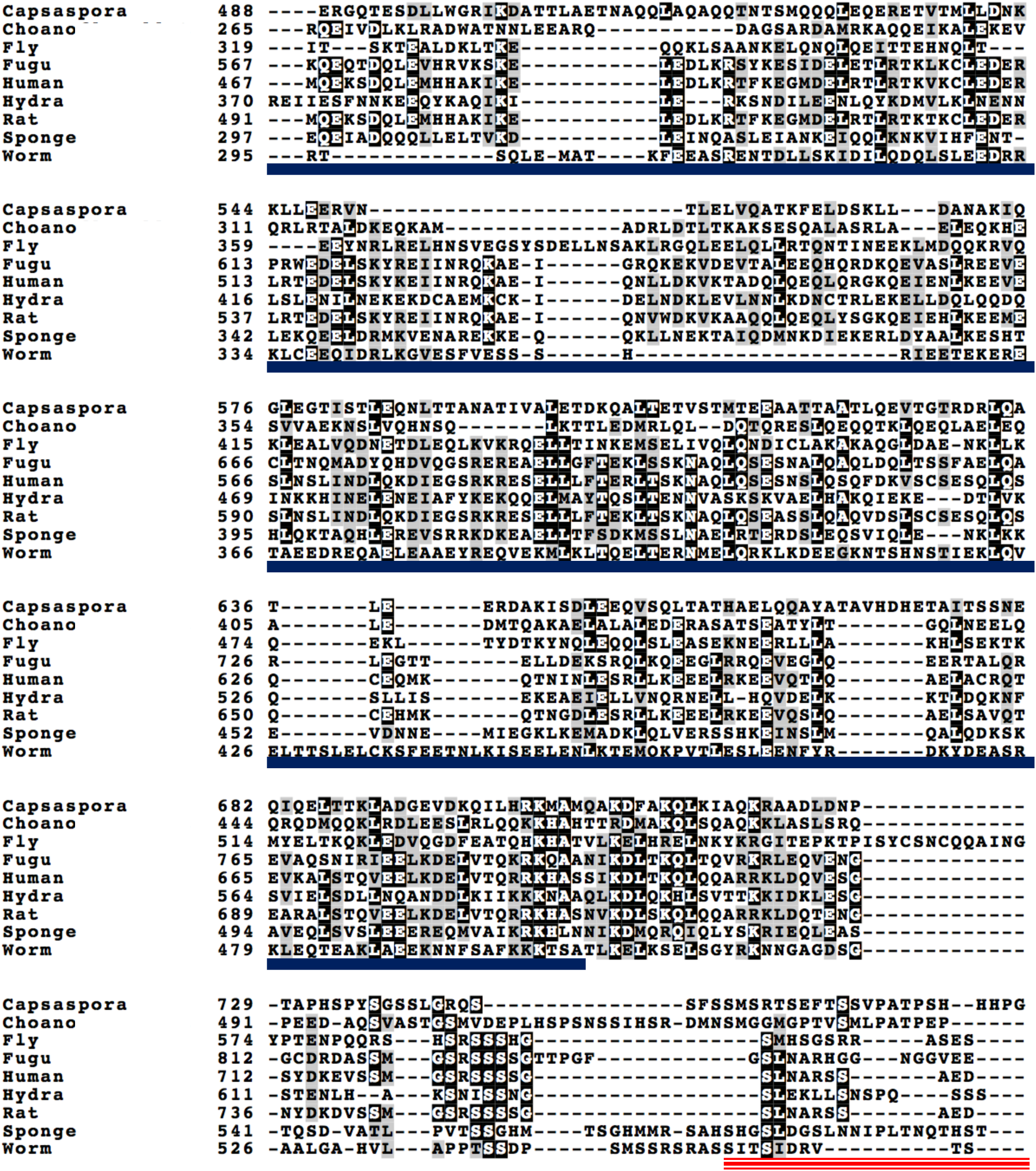
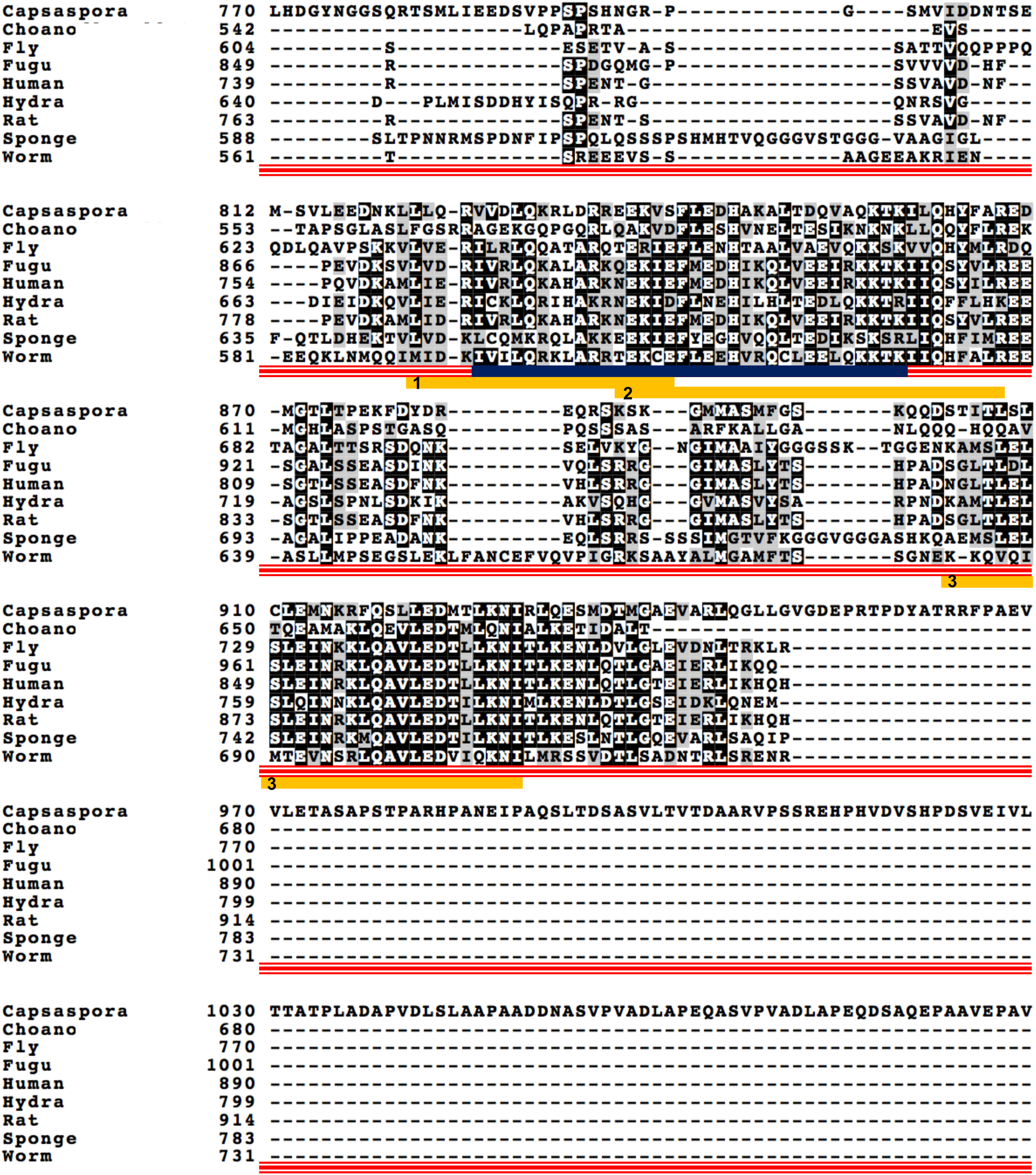
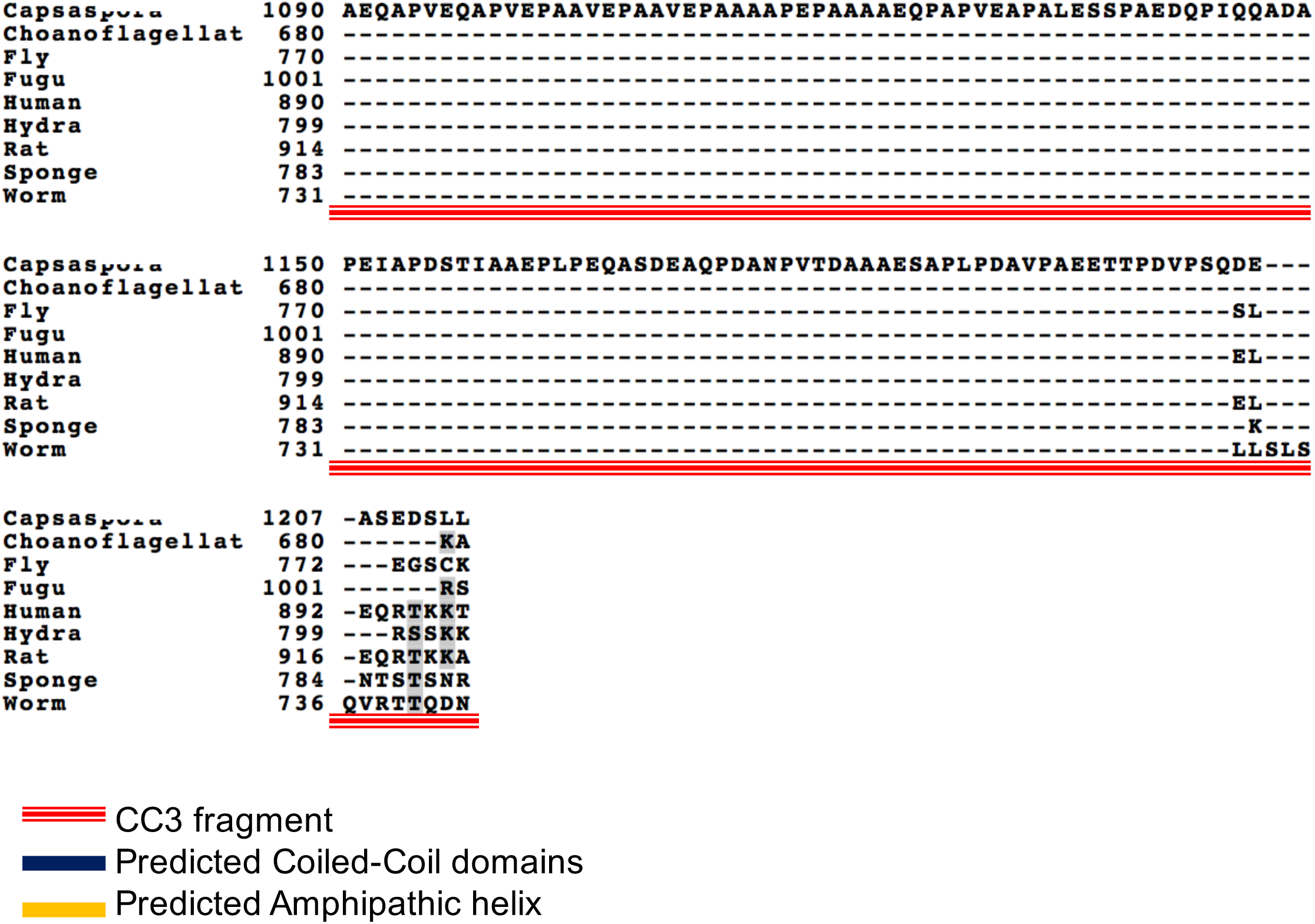
Alignment of the CCCP-1 protein. Alignment of CCCP-1 proteins from *Capsaspora owczarzaki* (Capsaspora, CAOG_00459, accession # XP_004365330.2), *Monosiga brevicollis* (Choano, hypothetical protein, accession # XP_001745351.1), *Drosophila melanogaster* (Fly, golgin 104, accession # NP_648879.1), *Takifugu rubripes* (Fugu, Ccdc186, XP_011604585.1), *Homo sapiens* (Human, Ccdc186, accession # AAI03500.1), *Hydra vulgaris* (Hydra, Ccdc186-like, XP_012558241.1), *Rattus norvegicus* (Rat, Ccdc186, accession # KX954625), *Amphimedon queenslandica* (Sponge, Ccdc186-like, XP_011409991.1), *C. elegans* (Worm, CCCP-1b, accession # NP_499628.1). Identical residues are shaded in black and similar residues are shaded in grey. Alignment was made with T-Coffee^1^, http://www.ebi.ac.uk/Tools/msa/tcoffee/, using default parameters and exhibited with BoxShade 3.21 (http://embnet.vital-it.ch/software/BOX_form.html). The coiled-coil domains of the worm protein (from SMART^2^, http://smart.embl-heidelberg.de) are marked with blue bars. The CC3 domain of worm CCCP-1 is marked with a red bar. The presence of potential amphipathic helices in the CC3 domain was determined using the following settings in HeliQuest^3^ (http://heliquest.ipmc.cnrs.fr): Helix type: ⍺, window size: 18 amino acids, Hydrophobic moment (µH) peaks above 0.35. The predicted helices with a patch of five or more hydrophobic amino acids are numbered 1, 2 and 3. In the human ortholog, the region of helices #1 and #2 is still predicted to form amphipathic helices, but the region of helix #3 is not predicted to form an amphipathic helix.

1. C N, Dg H, J H. T-Coffee: A novel method for fast and accurate multiple sequence alignment. *J Mol Biol*. 2000;302(1):205-217.
2. Schultz J, Milpetz F, Bork P, Ponting CP. SMART, a simple modular architecture research tool: Identification of signaling domains. *Proc Natl Acad Sci*. 1998;95(11):5857-5864.
3. Gautier R, Douguet D, Antonny B, Drin G. HELIQUEST: a web server to screen sequences with specific α-helical properties. *Bioinformatics*. 2008;24(18):2101-2102.

**Table S1.**
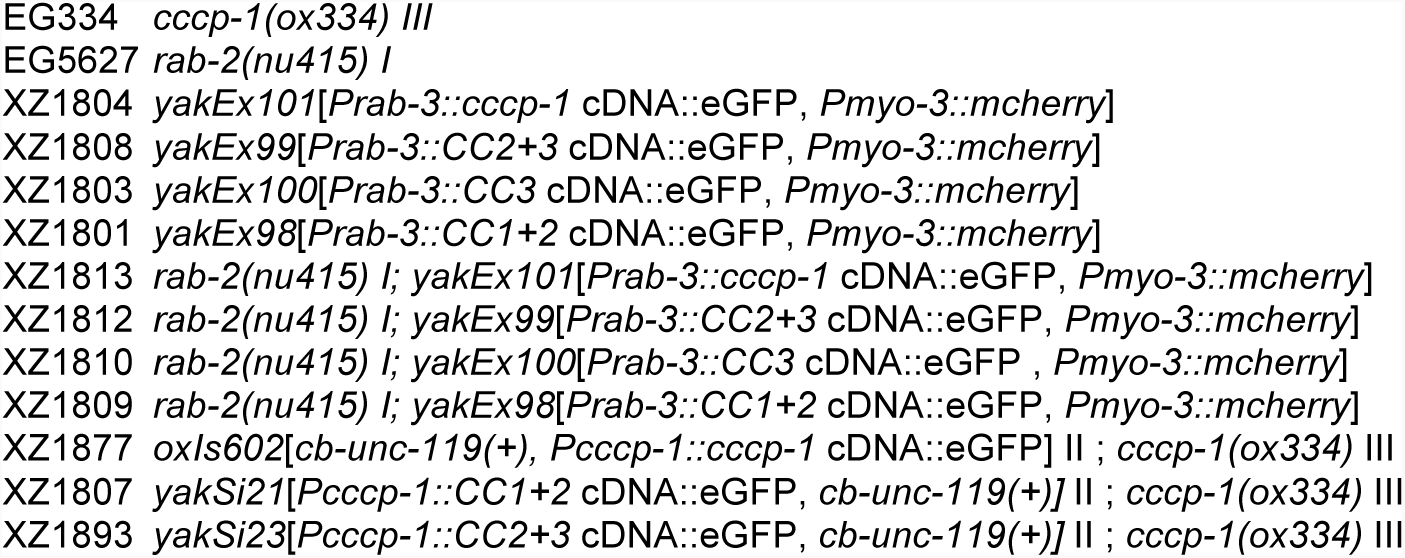
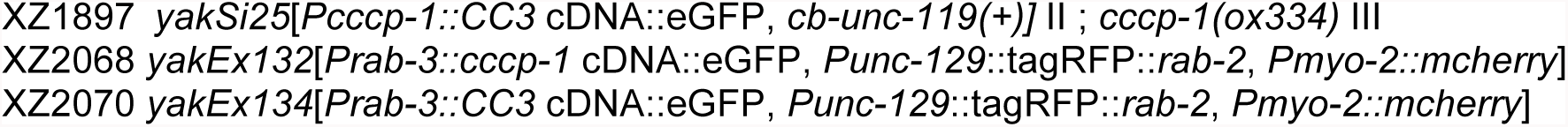
Strain list.

**Table S2.**
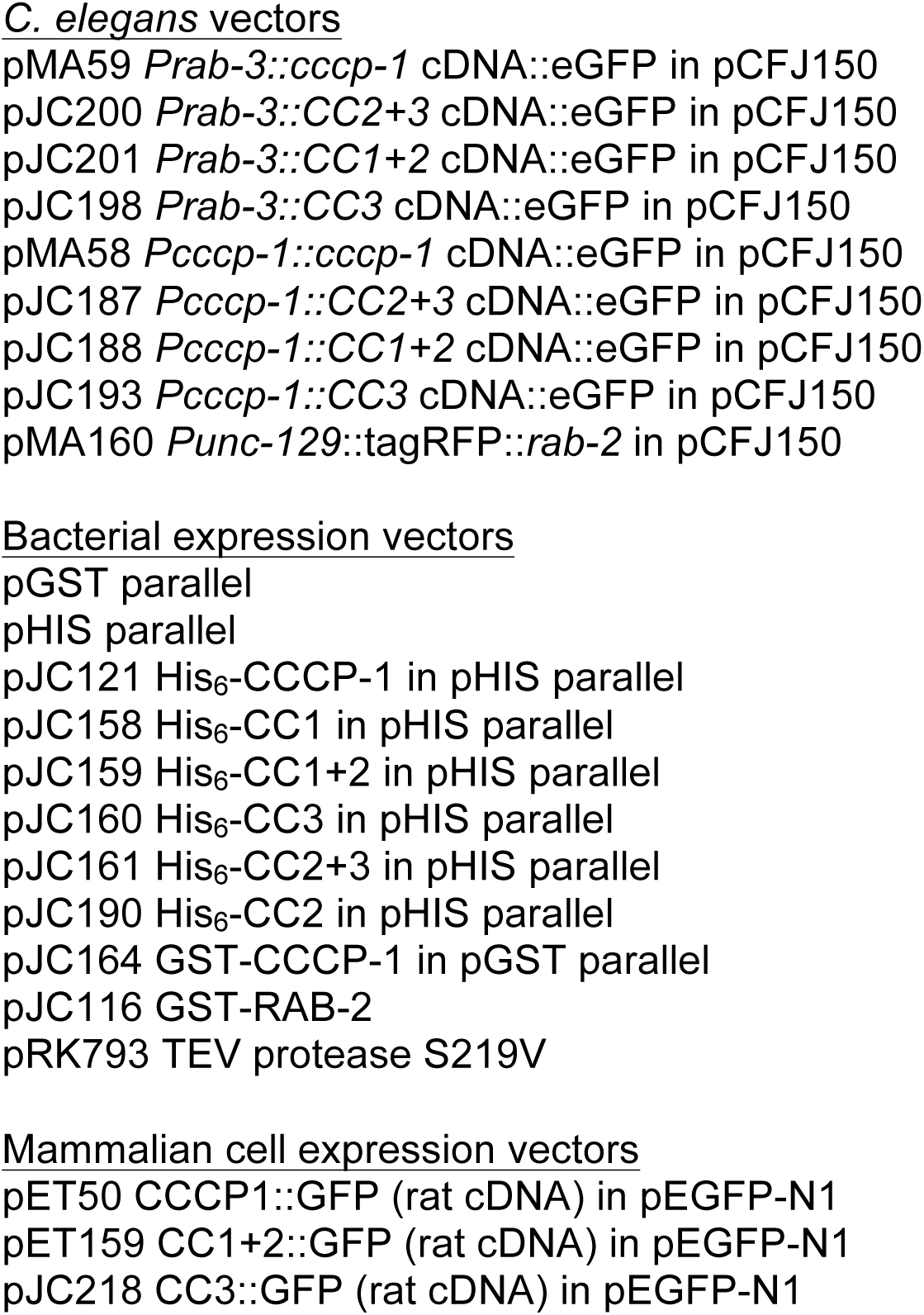
Plasmid list.

## References

1. Tooze SA, Martens GJ, Huttner WB. Secretory granule biogenesis: rafting to the SNARE. Trends Cell Biol. 2001;11(3):116–122.

2. Morvan J, Tooze SA. Discovery and progress in our understanding of the regulated secretory pathway in neuroendocrine cells. Histochem Cell Biol. 2008;129(3):243–252.

3. Borgonovo B, Ouwendijk J, Solimena M. Biogenesis of secretory granules. Curr Opin Cell Biol. 2006;18(4):365–370.

4. Gondré-Lewis MC, Park JJ, Loh YP. Cellular mechanisms for the biogenesis and transport of synaptic and dense-core vesicles. Int Rev Cell Mol Biol. 2012;299:27–115.

5. Bonnemaison M, Bäck N, Lin Y, Bonifacino JS, Mains R, Eipper B. AP-1A Controls Secretory Granule Biogenesis and Trafficking of Membrane Secretory Granule Proteins. Traffic. 2014;15(10):1099–1121.

6. Asensio CS, Sirkis DW, Edwards RH. RNAi screen identifies a role for adaptor protein AP-3 in sorting to the regulated secretory pathway. J Cell Biol. 2010;191(6):1173–1187.

7. Asensio CS, Sirkis DW, Maas Jr. JW, et al. Self-Assembly of VPS41 Promotes Sorting Required for Biogenesis of the Regulated Secretory Pathway. Dev Cell. 2013;27(4):425–437.

8. Walter AM, Kurps J, de Wit H, et al. The SNARE protein vti1a functions in dense-core vesicle biogenesis. EMBO J. 2014;33(15):1681–1697.

9. Buffa L, Fuchs E, Pietropaolo M, Barr F, Solimena M. ICA69 is a novel Rab2 effector regulating ER– Golgi trafficking in insulinoma cells. Eur J Cell Biol. 2008;87(4):197–209.

10. Cao M, Mao Z, Kam C, et al. PICK1 and ICA69 Control Insulin Granule Trafficking and Their Deficiencies Lead to Impaired Glucose Tolerance. PLoS Biol. 2013;11(4):e1001541.

11. Holst B, Madsen KL, Jansen AM, et al. PICK1 Deficiency Impairs Secretory Vesicle Biogenesis and Leads to Growth Retardation and Decreased Glucose Tolerance. PLoS Biol. 2013;11(4):e1001542.

12. Pinheiro PS, Jansen AM, de Wit H, et al. The BAR Domain Protein PICK1 Controls Vesicle Number and Size in Adrenal Chromaffin Cells. J Neurosci. 2014;34(32):10688–10700.

13. Kebede MA, Oler AT, Gregg T, et al. SORCS1 is necessary for normal insulin secretory granule biogenesis in metabolically stressed β cells. J Clin Invest. 2014;124(10):4240–4256.

14. Hao Z, Wei L, Feng Y, et al. Impaired maturation of large dense-core vesicles in muted-deficient adrenal chromaffin cells. J Cell Sci. 2015;128(7):1365–1374.

15. Torres IL, Rosa-Ferreira C, Munro S. The Arf family G protein Arl1 is required for secretory granule biogenesis in Drosophila. J Cell Sci. 2014;127(10):2151–2160.

16. Burgess J, Jauregui M, Tan J, et al. AP-1 and clathrin are essential for secretory granule biogenesis in Drosophila. Mol Biol Cell. 2011;22(12):2094–2105.

17. Ahras M, Otto GP, Tooze SA. Synaptotagmin IV is necessary for the maturation of secretory granules in PC12 cells. J Cell Biol. 2006;173(2):241–251.

18. Wendler F, Page L, Urbé S, Tooze SA. Homotypic Fusion of Immature Secretory Granules During Maturation Requires Syntaxin 6. Mol Biol Cell. 2001;12(6):1699–1709.

19. Edwards SL, Charlie NK, Richmond JE, Hegermann J, Eimer S, Miller KG. Impaired dense core vesicle maturation in Caenorhabditis elegans mutants lacking Rab2. J Cell Biol. 2009;186(6):881–895.

20. Sumakovic M, Hegermann J, Luo L, et al. UNC-108/RAB-2 and its effector RIC-19 are involved in dense core vesicle maturation in Caenorhabditis elegans. J Cell Biol. 2009;186(6):897–914.

21. Mesa R, Luo S, Hoover CM, et al. HID-1, a New Component of the Peptidergic Signaling Pathway. Genetics. 2011;187(2):467–483.

22. Hannemann M, Sasidharan N, Hegermann J, Kutscher LM, Koenig S, Eimer S. TBC-8, a Putative RAB-2 GAP, Regulates Dense Core Vesicle Maturation in Caenorhabditis elegans. PLoS Genet. 2012;8(5):e1002722.

23. Sasidharan N, Sumakovic M, Hannemann M, et al. RAB-5 and RAB-10 cooperate to regulate neuropeptide release in Caenorhabditis elegans. Proc Natl Acad Sci. 2012;109(46):18944–18949.

24. Ailion M, Hannemann M, Dalton S, et al. Two Rab2 Interactors Regulate Dense-Core Vesicle Maturation. Neuron. 2014;82(1):167–180.

25. Topalidou I, Cattin-Ortolá J, Pappas AL, Cooper K, Merrihew GE, MacCoss MJ, Ailion M. The EARP Complex and Its Interactor EIPR-1 Are Required for Cargo Sorting to Dense-Core Vesicles. PLOS Genet 2016;12:e1006074.

26. Stenmark H. Rab GTPases as coordinators of vesicle traffic. Nat Rev Mol Cell Biol. 2009;10(8):513–525.

27. Gillingham AK, Sinka R, Torres IL, Lilley KS, Munro S. Toward a Comprehensive Map of the Effectors of Rab GTPases. Dev Cell. 2014;31(3):358–373.

28. Gillingham AK, Munro S. Long coiled-coil proteins and membrane traffic. Biochim Biophys Acta BBA - Mol Cell Res. 2003;1641(2–3):71–85.

29. Gillingham AK, Munro S. Finding the Golgi: Golgin Coiled-Coil Proteins Show the Way. Trends Cell Biol. 2016;26(6):399–408.

30. Witkos TM, Lowe M. The Golgin Family of Coiled-Coil Tethering Proteins. Front Cell Dev Biol. 2016;3.

31. Yu I-M, Hughson FM. Tethering factors as organizers of intracellular vesicular traffic. Annu Rev Cell Dev Biol. 2010;26:137–156.

32. Sztul E, Lupashin V. Role of vesicle tethering factors in the ER-Golgi membrane traffic. FEBS Lett. 2009;583(23):3770–3783.

33. Hohmeier HE, Mulder H, Chen G, Henkel-Rieger R, Prentki M, Newgard CB. Isolation of INS-1-derived cell lines with robust ATP-sensitive K+ channel-dependent and -independent glucose-stimulated insulin secretion. Diabetes. 2000;49(3):424–430.

34. Bigay J, Casella J-F, Drin G, Mesmin B, Antonny B. ArfGAP1 responds to membrane curvature through the folding of a lipid packing sensor motif. EMBO J. 2005;24(13):2244–2253.

35. Sapperstein SK, Walter DM, Grosvenor AR, Heuser JE, Waters MG. p115 is a general vesicular transport factor related to the yeast endoplasmic reticulum to Golgi transport factor Uso1p. Proc Natl Acad Sci U S A. 1995;92(2):522–526.

36. Ishida R, Yamamoto A, Nakayama K, et al. GM130 is a parallel tetramer with a flexible rod-like structure and N–terminally open (Y-shaped) and closed (I-shaped) conformations. FEBS J. 2015;282(11):2232–2244.

37. Cheung PP, Limouse C, Mabuchi H, Pfeffer SR. Protein flexibility is required for vesicle tethering at the Golgi. eLife. 2016;4:e12790.

38. Munro S, Nichols BJ. The GRIP domain – a novel Golgi-targeting domain found in several coiled-coil proteins. Curr Biol. 1999;9(7):377–380.

39. Antonny B, Beraud-Dufour S, Chardin P, Chabre M. N-terminal hydrophobic residues of the G-protein ADP-ribosylation factor-1 insert into membrane phospholipids upon GDP to GTP exchange. Biochemistry (Mosc). 1997;36(15):4675–4684.

40. Drin G, Antonny B. Amphipathic helices and membrane curvature. FEBS Lett. 2010;584(9):1840–1847.

41. Miller MB, Vishwanatha KS, Mains RE, Eipper BA. An N-terminal Amphipathic Helix Binds Phosphoinositides and Enhances Kalirin Sec14 Domain-mediated Membrane Interactions. J Biol Chem. 2015;290(21):13541–13555.

42. Gautier R, Douguet D, Antonny B, Drin G. HELIQUEST: a web server to screen sequences with specific α-helical properties. Bioinformatics. 2008;24(18):2101–2102.

43. Drin G, Casella J-F, Gautier R, Boehmer T, Schwartz TU, Antonny B. A general amphipathic α-helical motif for sensing membrane curvature. Nat Struct Mol Biol. 2007;14(2):138–146.

44. Brown FC, Schindelhaim CH, Pfeffer SR. GCC185 plays independent roles in Golgi structure maintenance and AP-1–mediated vesicle tethering. J Cell Biol. 2011;194(5):779–787.

45. Wong M, Munro S. The specificity of vesicle traffic to the Golgi is encoded in the golgin coiled-coil proteins. Science. 2014;346(6209):1256898.

46. Urbé S, Page LJ, Tooze SA. Homotypic Fusion of Immature Secretory Granules during Maturation in a Cell-free Assay. J Cell Biol. 1998;143(7):1831–1844.

47. Du W, Zhou M, Zhao W, et al. HID-1 is required for homotypic fusion of immature secretory granules during maturation. eLife. 2016;5:e18134.

48. Ailion M, Thomas JH. Isolation and Characterization of High-Temperature-Induced Dauer Formation Mutants in Caenorhabditis elegans. Genetics. 2003;165(1):127–144.

49. Yu Y, Wang L, Jiu Y, et al. HID-1 is a novel player in the regulation of neuropeptide sorting. Biochem J. 2011;434(3):383–390.

50. Arvan P, Halban PA. Sorting Ourselves Out: Seeking Consensus on Trafficking in the Beta-Cell. Traffic. 2004;5(1):53–61.

51. Klumperman J, Kuliawat R, Griffith JM, Geuze HJ, Arvan P. Mannose 6-phosphate receptors are sorted from immature secretory granules via adaptor protein AP-1, clathrin, and syntaxin 6-positive vesicles. J Cell Biol. 1998;141(2):359–371.

52. Schindler C, Chen Y, Pu J, Guo X, Bonifacino JS. EARP is a multisubunit tethering complex involved in endocytic recycling. Nat Cell Biol. 2015;17(5):639–650.

53. Brenner S. The Genetics of Caenorhabditis Elegans. Genetics. 1974;77(1):71–94.

54. Frøkjær-Jensen C, Davis MW, Hopkins CE, et al. Single copy insertion of transgenes in C. elegans. Nat Genet. 2008;40(11):1375–1383.

55. Frøkjær-Jensen C, Davis MW, Ailion M, Jorgensen EM. Improved Mos1-mediated transgenesis in C. elegans. Nat Methods. 2012;9(2):117–118.

56. Gibson DG, Young L, Chuang R-Y, Venter JC, Hutchison CA, Smith HO. Enzymatic assembly of DNA molecules up to several hundred kilobases. Nat Methods. 2009;6(5):343–345.

57. Gibson DG, Young L, Chuang R-Y, Venter JC, Hutchison CA, Smith HO. Enzymatic assembly of DNA molecules up to several hundred kilobases. Nat Methods 2009;6:343–345.

58. Mello CC, Kramer JM, Stinchcomb D, Ambros V. Efficient gene transfer in C.elegans: extrachromosomal maintenance and integration of transforming sequences. EMBO J. 1991;10(12):3959–3970.

59. Sheffield P, Garrard S, Derewenda Z. Overcoming Expression and Purification Problems of RhoGDI Using a Family of “Parallel” Expression Vectors. Protein Expr Purif. 1999;15(1):34–39.

60. Tropea JE, Cherry S, Waugh DS. Expression and Purification of Soluble His_6_-Tagged TEV Protease [Internet]. In: Doyle SA, editor. High Throughput Protein Expression and Purification: Methods and Protocols. Totowa, NJ: Humana Press; 2009. p. 297–307.

